# Simulation of neurotrophin receptor transmembrane helix interactions reveals active states and distinct signaling mechanisms

**DOI:** 10.1101/2024.08.16.608237

**Authors:** Christina Athanasiou, Ainara Claveras Cabezudo, Alexandros Tsengenes, Rebecca C. Wade

## Abstract

Neurotrophin (NT) receptor signaling regulates neuronal survival, axonal and dendritic network maintenance, differentiation, and synaptic plasticity. Signaling is initiated by NT binding to the extracellular domain of NT receptor dimers, leading to activation of the receptor and signal propagation intracellularly. How this activating signal is mediated by the single-pass transmembrane (TM) helical domain of the receptor, and what the relation between domain sequence and signaling mechanism is, remains unclear. The structure and dynamics of the TM domain of the receptor dimers in the active and inactive states for intracellular signaling are still elusive, with NMR structures capturing only a single state. Here, we carried out unbiased and enhanced sampling molecular dynamics simulations of the TM domain dimers of the wild-type p75, TrkA and TrkB NT receptors and selected mutants in micelle and bilayer lipid environments at atomistic and coarse-grained levels of representation. The coarse-grained simulations enabled exploration of multiple states of the TM domain dimers and revealed the influence of the lipid environment on the TM helix arrangements. From the simulations, we identify active and inactive TM helix arrangements of the p75 and TrkA receptors that are supported by experimental data and suggest two different signaling mechanisms through the C-terminal regions of the TM helices. For TrkB, a single dominant but less energetically stable arrangement of the TM domain dimer is observed. These findings have implications for mechanistic studies of NT receptor signaling and the design of neuroprotective drugs to stabilize specific states of the TM domain of the receptors.

**Significance Statement:** Neurotrophins regulate neuronal survival, growth and differentiation during development, and play a role in many neurodegenerative and psychiatric disorders. They initiate signaling through the cell membrane by associating extracellularly with transmembrane receptors. The transmembrane helical domain of the neurotrophin receptors is responsible for transmitting the activation signal to the cell interior. However, how the transmembrane domain mediates signal transmission and the relation between its sequence and signaling mechanism remain unclear. Here, by employing state-of-the-art molecular dynamics simulation techniques, we identify active and inactive states of the transmembrane domains of the three main neurotrophin receptors that support distinct transmembrane signaling mechanisms for these receptors. Our results have implications for mechanistic studies of neurotrophin signaling and the design of neuroprotective drugs.

## INTRODUCTION

Neurotrophin (NT) proteins are members of a family of neurotrophic factors that have key roles in controlling the development and function of the central and peripheral nervous systems. [1], [2] The levels of secreted NTs modulate signaling pathways that regulate neuronal survival, axonal and dendritic network maintenance, differentiation, and synaptic plasticity.[3], [4] In addition to their physiological roles, NTs have been linked to neurodegenerative disorders, including Alzheimer’s, Huntington’s, and Parkinson’s diseases, amyotrophic lateral sclerosis (ALS, or Lou Gehrig’s disease), and peripheral neuropathy.[5], [6]

NTs exert their actions through binding to two different classes of single-pass transmembrane receptors: the tropomyosin receptor kinases A, B and C (TrkA, TrkB, TrkC), which are receptor tyrosine kinases, and the p75 receptor, which is a tumor necrosis factor receptor. The extracellular (EC) part of the Trks contains five domains: two cysteine-rich domains (D1, D3) flanking a leucine-rich domain (D2), and two immunoglobulin-like (Ig-like) domains (D4, D5). On the other hand, p75 has four cysteine-rich domains (CRD1-4) in the EC part (**Figure 1A**). Both types of receptor have a helical transmembrane (TM) domain connected via flexible linkers to the EC and the intracellular (IC) domains, with the latter being a kinase in the Trks and a death domain in p75. NTs bind to the EC segments of receptor dimers (**Figure 1A**) and thereby activate the receptors. The activation signal is transmitted to the IC domain through the TM helical domain, resulting in initiation of signaling inside the cell. Despite the TM domain being the focus of a number of in-depth studies,[7]–[9] the mechanisms by which the TM helices mediate receptor activation remain elusive.

**Figure 1:**
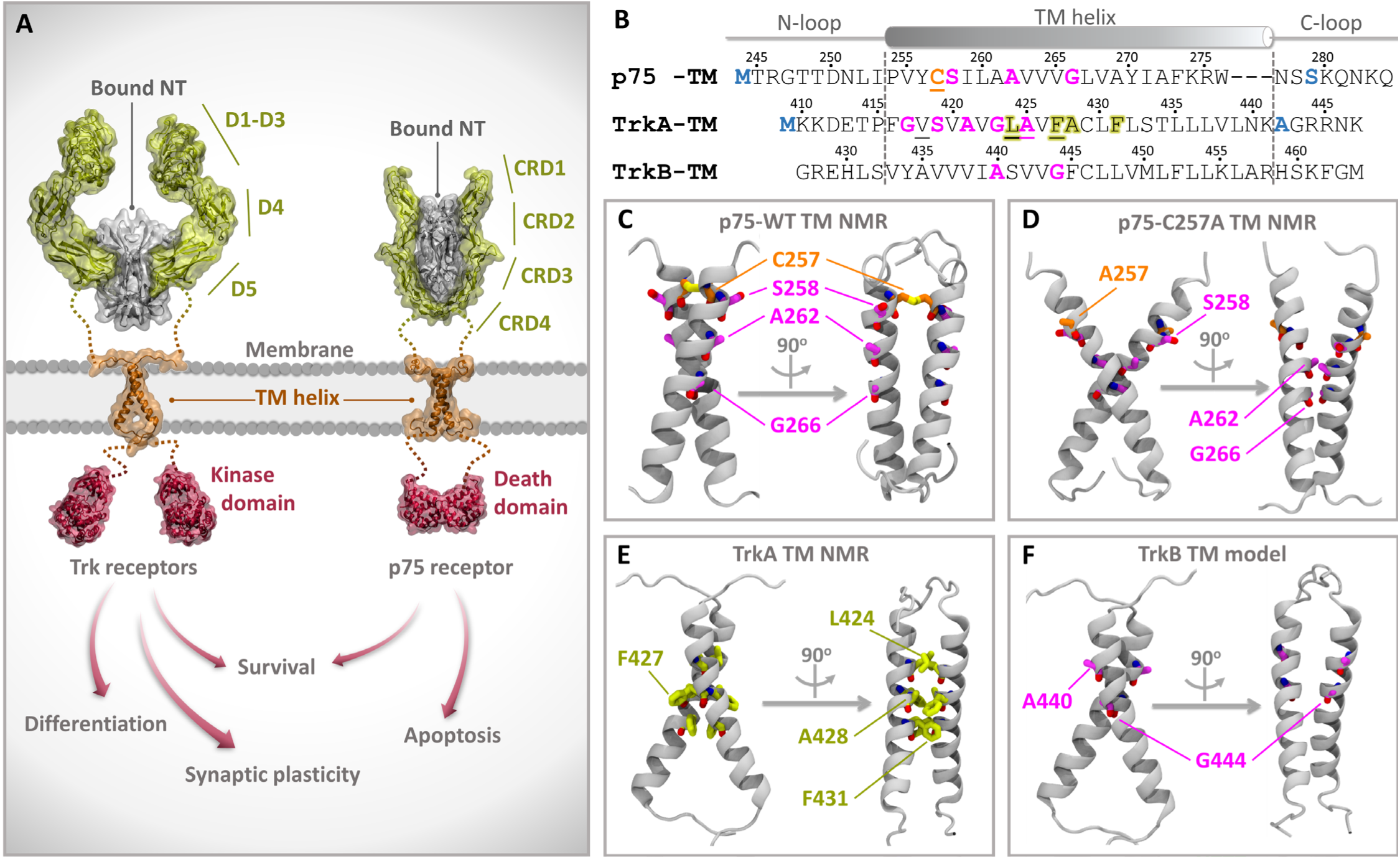
Sequence and structural features of NT receptors and their transmembrane (TM) domains. (A) Schematic representation of the Trk and p75 receptor homodimers with neurotrophin (NT) bound to the extracellular domains, indicating how NT binding leads to intracellular signal propagation. The domains whose structures have been determined experimentally are labelled and shown in cartoon representation with molecular surfaces whereas regions lacking experimentally determined structures are indicated by dotted lines. (B) Sequences used for the simulations of the TM domains of the receptors. The secondary structure, with the helical part indicated by dashed lines and flanked by N- and C-terminal loop regions, is shown as observed in the p75 (PDB ID: 2MIC)[9] and TrkA (PDB ID: 2N90)[7] NMR structures. The sequence of the human TrkB TM domain (Uniprot Q16620), for which we built a homology model, is shown aligned to TrkA. Mutated residues in the NMR structures compared to the human p75 (Uniprot P08138) and TrkA (Uniprot P04629) sequences are colored blue; the TM domains of rat (Uniprot P07174) and human p75 have the same sequence. The C257 residue in p75, which is mutated to Ala in the NMR structure of p75-C257A (PDB ID: 2MJO),[9] is underlined and colored orange. Residues participating in GxxxG-like motifs are colored magenta, while the TrkA L_424_xxF_427_A_428_xxF_431_ motif in the interface in the NMR structure is highlighted in yellow. (C-F) Cartoon representations of the (C) p75 WT (PDB ID: 2MIC), (D) p75-C257A (PDB ID: 2MJO), and (E) TrkA (PDB ID: 2N90) NMR structures, and the (F) homology model of the TrkB TM homodimer.

The predominant model for the activation of NT receptors requires dimerization even though it may also be possible for the receptors to act as monomers. [10], [11] Crosslinking, bimolecular fluorescence complementation and luciferase fragment complementation assays have shown that TrkA and TrkB can exist in the cell membrane prior to NT binding as preformed, inactive, homodimers, [12]–[14] or even higher oligomerization states. [15] Studies of truncation constructs showed that, in the absence of NT, dimerization of TrkA and TrkB is mediated by the TM and IC domains, [16] whereas p75 forms disulfide-linked homodimers through C257 in the TM domain, independently of NT binding. [17], [18] These results indicate that NT receptor dimers, and thus their respective TM domains, can exist in at least two states: active and inactive.

The TM helical dimer of p75 was the first to be structurally determined by Nuclear Magnetic Resonance (NMR) in dodecylphosphocholine (DPC) lipid micelles.[9] The p75 TM dimer is stabilized through a disulfide bridge at the C257 residue, independently of NT binding, with the C257A mutation abolishing NT-dependent receptor activity. [17], [18] The NMR structure of the p75 TM dimer in DPC micelles (**Figure 1B,C**) confirmed the existence of the disulfide bond, whereas the structure of the C257A mutant showed that its helices interact through a completely different interface that contains a GxxxG-like putative dimerization motif, A_262_xxxG_266_ (magenta residues in **Figure 1B,D**).[9]

Franco *et al.* recently solved the structure of the human TrkA TM domain homodimer in DPC lipid micelles by NMR, revealing an arrangement in which the TM helices interact through the L_424_xxF_427_A_428_xxF_431_ motif (yellow residues in **Figure 1B,E**).[7] Mutagenesis studies revealed that when TrkA is active, the TM monomers interact through a different interface with V418 playing a central role, suggesting that the NMR interface corresponds to the inactive state of the receptor.[7] Both active and inactive interfaces involve residues in the N-terminal half of the TM helix. The authors concluded that activation takes place through a rotation of the TM helices around their long axis.[7] Interestingly, the N-terminal TM region of the TrkA sequence contains three GxxxG-like putative dimerization motifs, G_417_xxxA_421_, S_419_xxxG_423_, and A_421_xxxA_425_, which might facilitate this rotation (magenta residues in **Figure 1B**). A combination of NMR, Förster Resonance Energy Transfer (FRET), molecular dynamics (MD) simulations and functional studies showed that, when the TM domains of TrkA and p75 monomers interact to form a heterodimer, they engage mainly through an interface that is opposite to the active interface and partially covering the inactive dimer interface, leaving the active interface exposed for binding to a second TrkA TM monomer.[19] Thus, binding of p75 to TrkA may favor the formation or stabilization of TrkA active homodimers. Furthermore, the extracellular linker of TrkA that connects the EC domains with the TM helix has been shown to play a key role in coupling the EC and TM segments upon activation, possibly through interacting with neurotrophic growth factor (NGF).[20]

A putative dimerization motif is also present in the TM sequence of human TrkB; A_440_xxxG_444_ (magenta residues in **Figure 1B,F**). Casarotto *et al.* recently found that, in MD simulations, the TrkB TM domain forms stable TM dimers that interact via the A_440_xxxG_444_ motif (A_439_xxxG_443_ in the rat TrkB sequence studied).[8] In this arrangement, the TrkB TM helices were observed to form a binding site for the antidepressant drug fluoxetine, which acts as a TrkB agonist. [8] Psychedelic drugs were also found to bind to a distinct but similar binding site at the interface of the TrkB TM helices, stabilizing the helices in a configuration favorable for activation.[21]

Here, to investigate the mechanisms by which the TM domain dimers mediate signal transduction to the cell interior, we explore the structural landscape of the TM domain dimers of the p75, TrkA and TrkB receptors by all-atom (AA) and coarse-grained (CG) MD simulations, using both unbiased and enhanced sampling techniques. We first assessed the ability of the AA and CG models to reproduce experimentally determined structures and to sample the biologically relevant states of the investigated systems. We then employed the Martini 2.2 [22], [23] and Martini 3[24] CG force fields, which have previously been successfully used to study similar systems, [25]–[29] for CG MD simulations of the TM domains in different lipid environments. In addition to conventional MD simulations, we performed metadynamics simulations to compute the free energy surfaces for TM helix dimerization. Analysis of the simulations provides insights into the signaling mechanisms of the TM domains of NT receptors and how, despite structural similarity, they differ between receptors, due to differences in the TM domain sequences.

## RESULTS

### CG MD simulations in a DPC micelle environment not only reproduce NMR structures but also explore other structural arrangements of the TM domain dimers

The NMR structures of the TM helix homodimers of WT p75,[9] p75 with the C257A mutation (p75-C257A),[9] and TrkA [7] have been determined in DPC micelles. Therefore, to first assess the ability of the simulations to reproduce experimental structures and sample possible TM helix dimer arrangements, we performed simulations of these TM helical dimers, and of a homology model of the TrkB TM domain dimer, in the same micelle environment, in atomistic and CG detail (**Figure 2A**). First, ten replica 1μs-MD simulations of the TrkA and TrkB TM helix homodimers were run with the Charmm36m all-atom (AA) force field [30] **(Table S1**, Systems 1 and 2). The TrkA and TrkB systems explore only left-handed helical arrangements (with positive crossing angles, Ω, **Figure 2B**) in the vicinity of the initial NMR and model structures, respectively, but not further, which suggests incomplete sampling at this atomistic level of representation (**Figure 2C, D**). Therefore, we next investigated the systems with CG simulations using the Martini 2.2 [22], [23] and Martini 3[24] force fields (**Table S1**, Systems 3-16). Ten replica CG simulations of 20 μs each were run for each system, resulting in a total simulation time of 1.6 ms. For the simulations of the helix homodimers in DPC micelles using the Martini 3 force field, we employed a scaling of the protein-water Lennard-Jones interactions, which we have previously shown to be necessary for the protein to be encapsulated into the micelle during the equilibration phase.[31] In contrast to the AA simulations, the CG simulations showed better exploration of the TM helix dimer arrangements, and since they are more computationally efficient, we also employed them for the simulation of the p75 and p75-C257A TM domain dimers (**Figure 2E-L**). The simulations of p75 with Martini 2.2 (**Figure 2E**) and Martini 3 (**Figure 2I**) force fields showed that the system explores configurations very similar to or the same as the NMR structure. This stability is conferred by the disulfide bond that connects the two helices of p75. In the absence of this disulfide bond in the p75-C257A mutant (**Figure 2F, J**), the system explores additional arrangements of the helices, with the Martini 3.3 simulations sampling very high crossing angles (Ω < -100° and Ω > 100°), which would probably not occur in the native environment of the cell membrane. These high crossing angles were observed in all simulations of p75-C257A, TrkA and TrkB in micelles using the Martini 3 force field.

**Figure 2:**
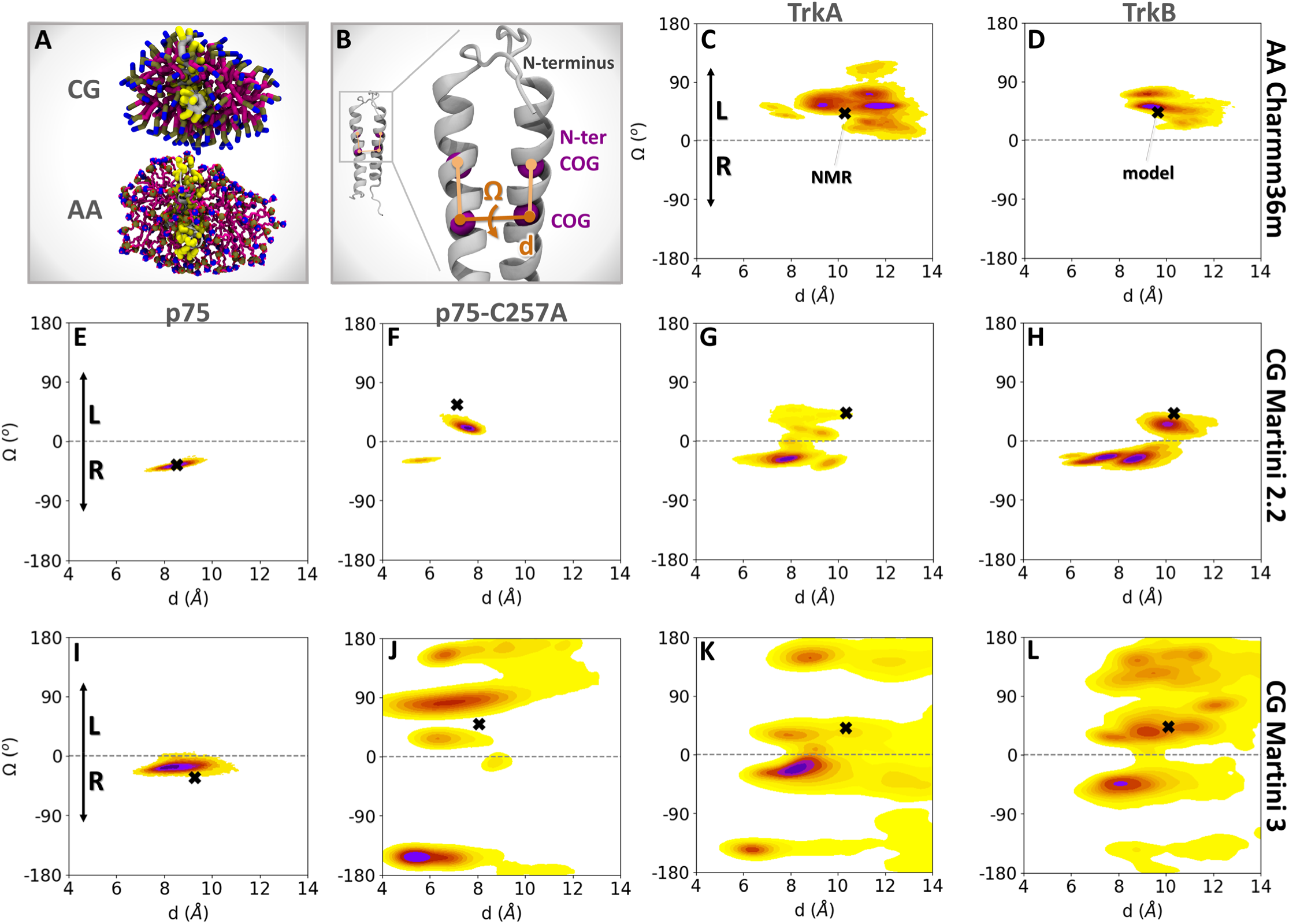
CG MD Simulations in a DPC micelle environment sample not only the NMR structures but also other low-energy arrangements of the two helices in the TM domain dimers. (A) Final snapshot of an equilibrated micelle formed by self-assembled DPC lipids (magenta) around a TrkA TM domain dimer (gray backbone with yellow side chains) extracted from one of the coarse-grained (CG) replica simulations. The system is shown in CG and all-atom (AA) representations. (B) Cartoon representation of a TM domain homodimer showing the definitions of the distance (d) between the centers of geometry (COG) of the backbone (BB) beads of the two helices and the crossing angle (Ω). The dihedral angle Ω is defined by the two planes formed by the COGs of the BB beads of the helices with the COGs of the BB beads of the N-terminal half of each of helix (N-ter COG). (C-L) Population density maps in the d-Ω space from (C,D) AA simulations with the Charmm36m force field, and CG simulations with the (E-H) Martini 2.2 and (I-L) Martini 3 force fields for p75, p75-C257A, TrkA and TrkB TM domain dimers in DPC micelles. The maps were computed by kernel density estimation (KDE) and the probability maxima are denoted in dark purple, with the color fading to yellow for states of lower probability. The NMR structures for p75, p75-C257A and TrkA and the homology model of TrkB, from which the simulations were started, are indicated with black crosses (**X**). Positive Ω values indicate left-handed helical dimers (L), while negative values correspond to right-handed dimers (R).

The TrkA and TrkB TM domain homodimers adopt arrangements close to the initial left-handed NMR and model structures, respectively, but also sample right-handed arrangements with both Martini 2.2 and Martini 3 force fields. Similar orientations of the helices were sampled with Martini 2.2 and 3, although there was more exploration in Martini 3. Interestingly, the TrkB homodimer was more stable than TrkA in the arrangement of the initial model (high sampling close to the black cross in **Figure 2H**), even though it was a homology model based on the NMR structure of TrkA. Additional test simulations with different equilibration protocols or conditions led to similar results (**Table S1**, Systems 7-12), indicating that the results are robust and not affected by these changes in simulation conditions.

Overall, the results confirm the ability of CG MD simulations to reproduce the NMR structures. All the receptors sample arrangements close to the initial NMR or model structures, respectively, with TrkA and TrkB also sampling additional arrangements of the two helices. For p75-C257A, the NMR structure appears more unstable than for WT p75, as would be expected. As Martini 3 offered more extensive sampling, probably due to the weaker nature of its protein-protein interactions compared to Martini 2.2,[32], we decided to employ Martini 3 for subsequent CG MD simulations.

### Simulations in a phospholipid bilayer environment capture NMR structures and reveal other distinct configurations of the TM domain dimers

The configurational landscape of TM helix homodimers in phospholipid bilayers is much more confined than that observed in micelles.[29] We attribute these differences to both the approximately planar geometry and to the less dynamic environment of bilayers, which contrast with the inherently dynamic and rather disordered nature of micelles. Extreme crossing angles higher than 100° or lower than -100°, as observed in the simulations of TM domain homodimers in micelles, would not be compatible with the bilayer thickness. To investigate the TM domain homodimers in the more realistic lipid environment of bilayers, CG simulations of the TM domain homodimers were run in 1-palmitoyl-2-oleoyl-sn-glycero-3-phosphocholine (POPC) membranes using the Martini 3 force field. For each receptor, 10 simulations were run, each of 20 μs duration (**Table S1**, Systems 17-20).

As in the micelle simulations, the TM domain dimer interface of p75 is highly stable due to the disulfide bridge at residue C257. This is well reflected in the 2D population density map (**Figure 3A**), which display a single maximum containing right-handed helical arrangements with a crossing angle Ω between -30° and -20° and a distance between centers of mass (COG) of the backbone beads (BB) of the helices, d, of ∼7.5 Å. In contrast, left- and right-handed helix dimers were observed in the simulations of p75-C257A, TrkA and TrkB (**Figure 3B-D**).

**Figure 3:**
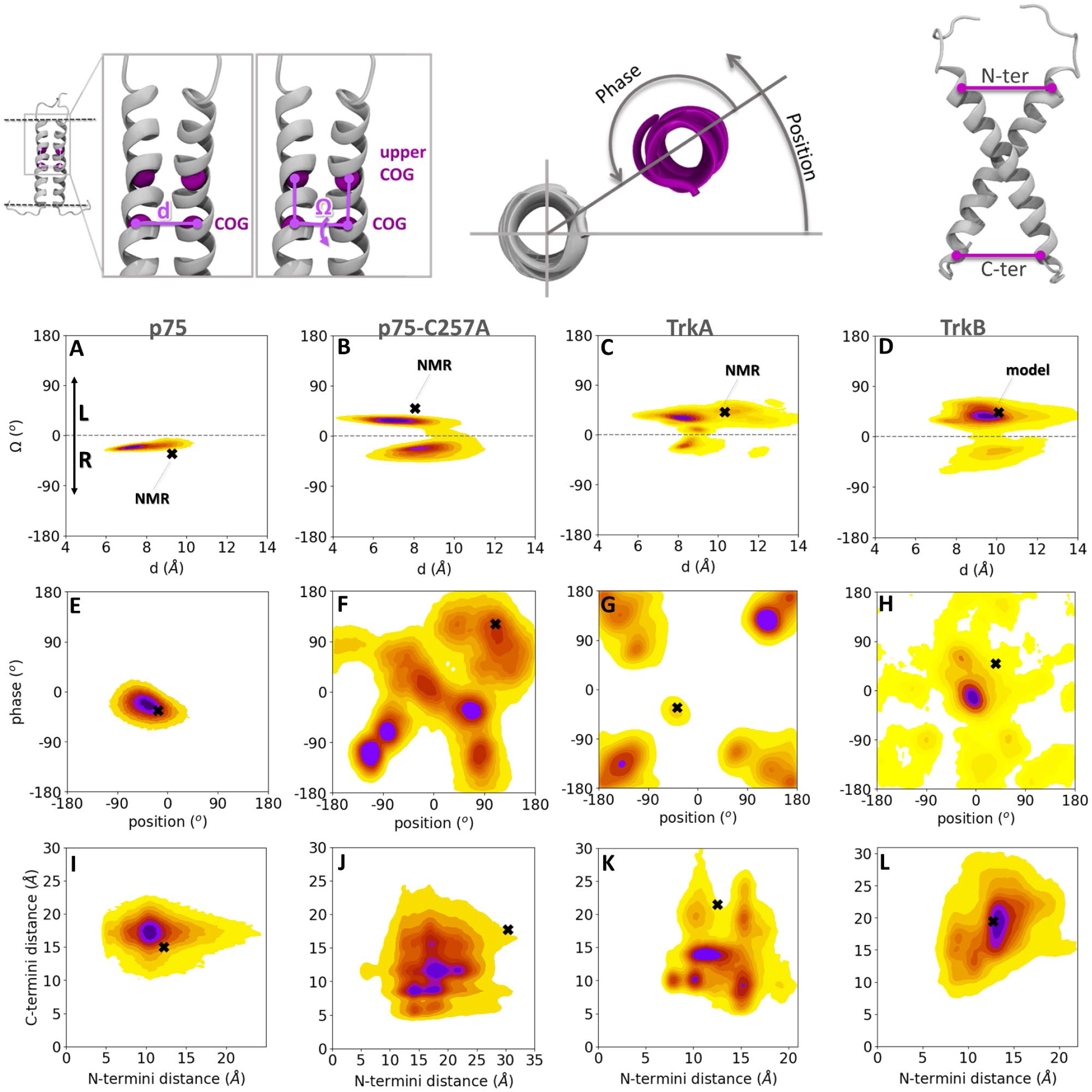
Simulations in a phospholipid bilayer environment show that the TM domain homodimers sample NMR structures and reveal other sequence-dependent low-energy arrangements of the two helices. Population density heatmaps from CG simulations with the Martini 3 force field of p75, p75-C257A, TrkA and TrkB in a POPC bilayer projected onto the 2D spaces of (A-D) d-Ω, (E-H) phase-position (the reference residues used for the phase-position calculations are V265 for p75 and p75-C257A, A428 for TrkA, and G444 for TrkB), and (I-L) C-termini and N-termini distances. The definitions of these geometric parameters are shown at the top of the figure. Colors and symbols as in Figure 2.

We next analyzed the torsion angles of the second helix with respect to the first helix and with respect to itself, referred to as position and phase, respectively [28] (see Materials and Methods section and **Figure 3**). Consistent with the stability of its interface, p75 exhibited minimal variation in both position and phase angles, (**Figure 3E, Figure S1**). The NMR structure of the p75 dimer is stabilized not only by the C257-C257 disulfide bond, but also by van der Waals interactions of L260, V264 and V265 and π-stacking of F273 and W276.[9] These interactions were preserved during the simulations as shown by the contact map between the residues of the two helices (**Figure S1**).

The C257A mutation in p75 leads to a destabilization of the dimer interface, which is manifested by several maxima in the phase-position density map (**Figure 3F**, **Figure S2**). The NMR structure of p75-C257A has the A262xxxG266 dimerization motif at the interface of the helices.[9] Analysis of the contacts between the TM helices in the various maxima shows that the dimerization motif is at least partially present in the interface in all maxima except the first, although the other contacts vary between the six different maxima (**Figure S2**).

As for p75-C257A, the TrkA helix dimers adopted a variety of arrangements during simulations, including the NMR structure (black cross in **Figure 3C,G**). The arrangements similar to the NMR structure correspond to a maximum in the phase-position density map, but interestingly, this maximum is not as highly populated in the simulations as other maxima, such as the most populated density maximum corresponding to dimers with rotation angles between 100° and 150° around both helices. In the simulations of TrkB, a dominant density maximum was observed with position and phase values close to 0°, which indicates that G444, which participates in the A440xxxG444 dimerization motif (**Figure 1B,F**) and which was used as the reference residue, is at the interface during most of the simulation time (**Figure 3H**, **Figure S1**; note that different residues were used as reference for the phase-position heatmaps for p75, TrkA and TrkB and thus, these heatmaps cannot be directly compared between the receptors). A high occupancy of contacts in the A440xxxG444 motif is also apparent in the contact map of the density maximum (**Figure S1**). This observation is in agreement with previous work, which has shown that the TrkB homodimer can be stabilized with this dimerization motif in an arrangement that probably corresponds to the active state of the receptor.[8] Additionally, S441 is at the interface of the helices with 100% occupancy in the dominant maximum configurations. This residue has been recently identified to mediate TrkB signaling and be at the interface of an NMR structure of the TrkB TM homodimer that was determined after our simulations were carried out. [33]

The distances between the helix termini were also monitored. In agreement with the other descriptors of the configurational landscape, the simulations of the p75-C257A and TrkA dimers revealed several density maxima with distinct combinations of N- and C-termini distances, in contrast to p75 and TrkB, for which a single density maximum was observed (**Figure 3I-L**).

### Active and inactive configurations of the TrkA-TM domain dimer are identified in the CG MD simulations

As discussed above, mutagenesis studies [7] indicate that there are two interfaces of the TM helices in TrkA, an active and an inactive one, with the solved NMR structure corresponding to the inactive interface as shown in **Figure 4A**. Surprisingly, considering that the simulations started from the NMR structure, the dominant maximum population density configuration in the phase-position distribution from the simulations, with rotation angles between 100° and 150°, has interfacial interactions involving residues (V422, A425, C429 and L432) that were suggested from the mutagenesis studies to be part of the active interface (**Figure 4A,B,C**). This finding suggests that the CG simulations of TrkA predominantly sample an arrangement of the TM helices that corresponds to the physiologically active state of the receptor.

**Figure 4:**
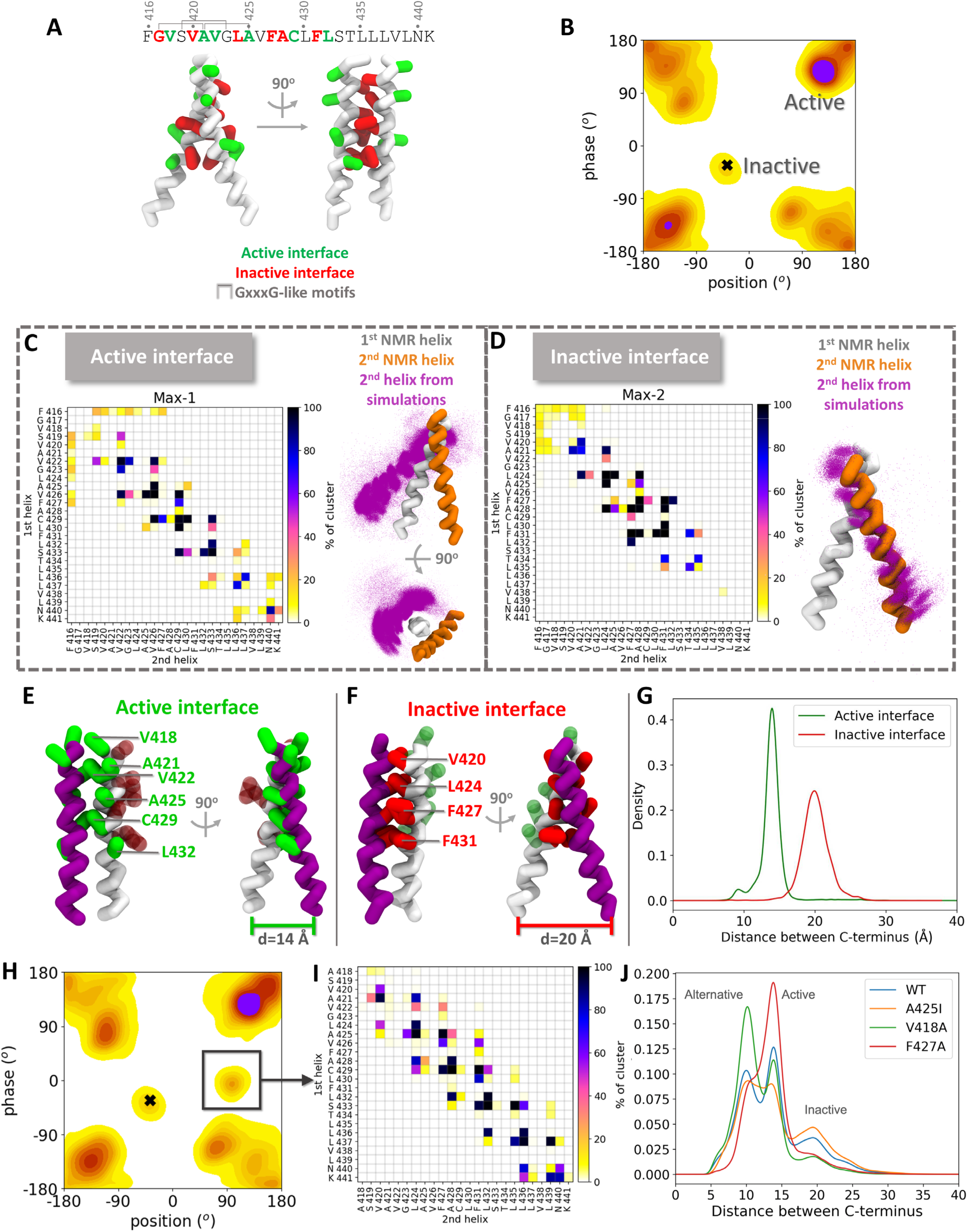
Comparison of active and inactive arrangements of the TrkA TM domain dimer in the CG MD simulations reveals closer C-terminii in the active arrangement. (A) NMR structure of the TrkA TM domain (PDB ID: 2N90) [7] in CG representation in an inactive arrangement with the residues of the active and inactive interfaces (as defined by mutagenesis data in [7]) shown in green and red, respectively. The three GxxxG-like motifs are indicated in the sequence. (B) Phase-position population density map from the CG Martini 3 simulations, with the two maxima that correspond to the active and inactive arrangements labelled. (C, D) Contact maps corresponding to the (C) active and (D) inactive TrkA arrangements labelled in (B). A contact is defined when a CG bead from one helix is within 6 Å of a bead of the second helix. The corresponding TM domain arrangements are shown with all the frames aligned to the 1^st^ helix (white). The 2^nd^ helix from the NMR structure (with an inactive arrangement) is shown in orange for reference, and the 2^nd^ helix from the simulation frames is shown in purple dots. Only backbone beads are depicted for simplicity. (E, F) Interface residues in the simulations in the (E) active and (F) inactive arrangements from representative frames of the active and inactive maxima in (B). (G) Population density plot from the simulations of the distance between the C-termini of the helices from the active and inactive maxima in (B). (H) Phase-position population density map of the TrkA V418A mutant and (I) corresponding contact map between the two helices in the indicated emergent region. (J) Population density plot of the distance between the C-termini for the WT TrkA TM domain and selected mutants. See Fig. S3 and S4 for further population density heatmaps and for helix contact difference maps, respectively, for these and other TrkA mutants.

In comparison, the contact map of the peak in the phase-position distribution that corresponds to the NMR structure (black cross, **Figure 4B**) shows the L_424_xxF_427_A_428_xxF_431_ motif at the interface (**Figure 4D**). This motif is present in the NMR structure and has been suggested to be part of the inactive interface (**Figure 4A**). Comparison of the TM helix arrangements from the active and inactive interfaces in the simulations and the NMR structure (**Figure 4C**,**D**) shows that the active interface is located on the opposite side of the helix to the inactive NMR interface, as was suggested from mutagenesis data.[7]

The proximity of the C-terminal kinase domains of TrkA is essential for autophosphorylation and subsequent signal propagation.[34] Hence, we monitored the C-terminal interhelical distance during the simulations. The distance between C-termini is shorter at ∼14 Å in the active state than in the inactive state where it is ∼20 Å (**Figure 4E,F,G**). This difference in the distance between C-termini may relate to receptor function: the approach of the two TM helix C-termini to each other could lead to propagation of the activation signal internally to the intracellular domains.

### TrkA TM domain mutations affect active and inactive interfaces

To assess the importance of specific contacts in both the active and inactive interfaces of TrkA, we generated single point mutations at key residues and subsequently ran simulations of the mutant TM domain dimers in POPC bilayers (**Table S1,** Systems 21-26). Hydrophobic residues (leucine, phenylalanine and valine) were mutated to alanine to study the effect of side-chain removal without compromising the helical propensity of the TM domain. In addition, following Franco and co-authors, who found that mutations at the active interface led to a drop in functional activity, the small residue alanine was mutated to the bulkier isoleucine, with the aim of inducing steric clashes and thereby disrupting the dimer interface.[7]

While the overall structural landscapes in the mutated systems appear similar to the wild-type system, analysis of the structural ensembles revealed systematic sequence dependent differences in the dimer interfaces, as well as shifts in the relative populations of the configurations (**Figure S3**). For the V418A mutant, for instance, right-handed helical dimer arrangements with negative crossing angles were more frequent than for WT TrkA (**Figure S3A**). In the simulations of this mutant, a minor population was sampled in a region of the configurational space not populated by WT TrkA with position and phase angles around 90° and 0°, respectively (**Figure 4H**). This arrangement has a heterotypic interface, containing residues from the active and inactive interfaces in different helices (**Figure 4I**). Although the interfacial interactions of V418 are weak and hardly change upon mutation to alanine, differences in the occurrence of inter-helical contacts with respect to the WT receptor were observed, most of them resulting in a shift of the interface by one residue towards the N-terminus. For instance, the contacts of V426 and L430 were replaced by the interactions of A425 and C429, respectively (**Figure S4A**).

It was previously reported that the V418C mutation induced dimerization via the formation of a disulfide bond and subsequent activation of TrkA.[7] In our simulations (without a disulfide bond at residue 418), this mutant did not show major differences with respect to WT TrkA in terms of the sampled conformational landscape (**Figure S3D-F**). The additional minor region described above in the conformational landscape of the V418A mutant was not observed for the V418C mutant **(Figure S3E**). However, in the active state, the two C418 residues from the two helices are in close proximity, and thus would be primed to form a covalent bond. Consistent with the formation of a disulfide bond, an increase in contacts between C418 residues was observed with respect to the WT simulations (**Figure S4B**).

The mutation of F427 to alanine induced the greatest changes in the configurational landscape and the contact interface of the TrkA TM domain dimer. Configurations with rotation angles close to those observed in the NMR structure were not observed for the F427A TrkA mutant (**Figure S3Q**) (or, indeed, for the L424A mutant (**Figure S3K**)). Furthermore, the dominant arrangement shifted to one with negative position angles. In contrast to the V418 mutants, which mainly showed contact changes in the N-terminal region, interactions along the whole length of the helices of the F427A mutant were perturbed (**Figure S4F**).

We identified three density peaks that were consistently present at identical C-termini distances across all systems, but with differences in their relative occurrence. We attribute the peak with the shortest distance to alternative arrangements to the active and inactive ones, as it does not overlap with the distance peaks observed for the WT active and inactive arrangements described above (**Figure 4J** vs **4G**). The V418A mutation resulted in an increase of arrangements with this very short distance between C-termini and intermediate interfaces. In agreement with the hypothesis that F427 participates in the inactive interface, mutation of this residue induced a decrease in inactive arrangements and an increase in active arrangements with C-terminal distances around 14 Å. The opposite effect is observed for the A425I mutant, consistent with a potential role of this residue in the active interface, as was proposed by Franco et al.[7].

### The free energy surface for TM domain dimerization indicates lower stability of the TrkB TM helix homodimer

CG-metadynamics (CG-MetaD) simulations were carried out to compute the underlying free energy surface (FES) for helix dimerization in the p75-C257A, TrkA and TrkB systems (**Table S1**, Systems 27-32). This enhanced sampling simulation protocol combines CG simulations with well-tempered metadynamics and it has been successfully used to study epidermal growth factor receptor (EGFR) TM helix dimerization.[25] The CG-MetaD simulations were run with both the Martini 2.2 and Martini 3 force fields, reaching a cumulative simulation time of ∼ 760 μs for all systems. The interhelical distance (d) was defined as the biasing collective variable (CV) in the simulations to allow exploration of a wide range of distances, thus enabling sampling of multiple helix-helix association and dissociation events and the calculation of the free energy of TM helix dimerization, ΔG_bind,_ from the FES as the difference between the bound and unbound states see **Figure 5A**. Convergence of the CG-MetaD simulations was verified (**Figures S5, S6**). More favorable binding free energies of the helices were computed with Martini 2.2 than Martini 3, in agreement with previous observations of the more attractive nature of protein-protein interactions in the Martini 2.2 force field,[32]. The binding free energies of the p75-C257A and TrkA TM helices measured by titration in DPC micelles are very similar at -7.5 kJ/mol and -7.9 ± 0.8 kJ/mol, respectively.[7], [9] Although a direct comparison of these values with the computed values for ΔG_bind_ in a POPC bilayer is not possible, the values for ΔG_bind_ computed with Martini 3 are closer than those from Martini 2, and also consistently, the computed ΔG_bind_ for TrkA is similar to that for p75-C257A in the Martini 3 simulations. In contrast, the dimerization free energy of TrkB is computed to be approximately half that of the other two receptors. For TrkB, ΔG_bind_ = -11 ± 0.35 kJ/mol, compared to ΔG_bind_ = -26 ± 0.53 kJ/mol for TrkA, and ΔG_bind_ = -19 ± 0.46 kJ/mol for p75-C257A, in a POPC bilayer.

**Figure 5:**
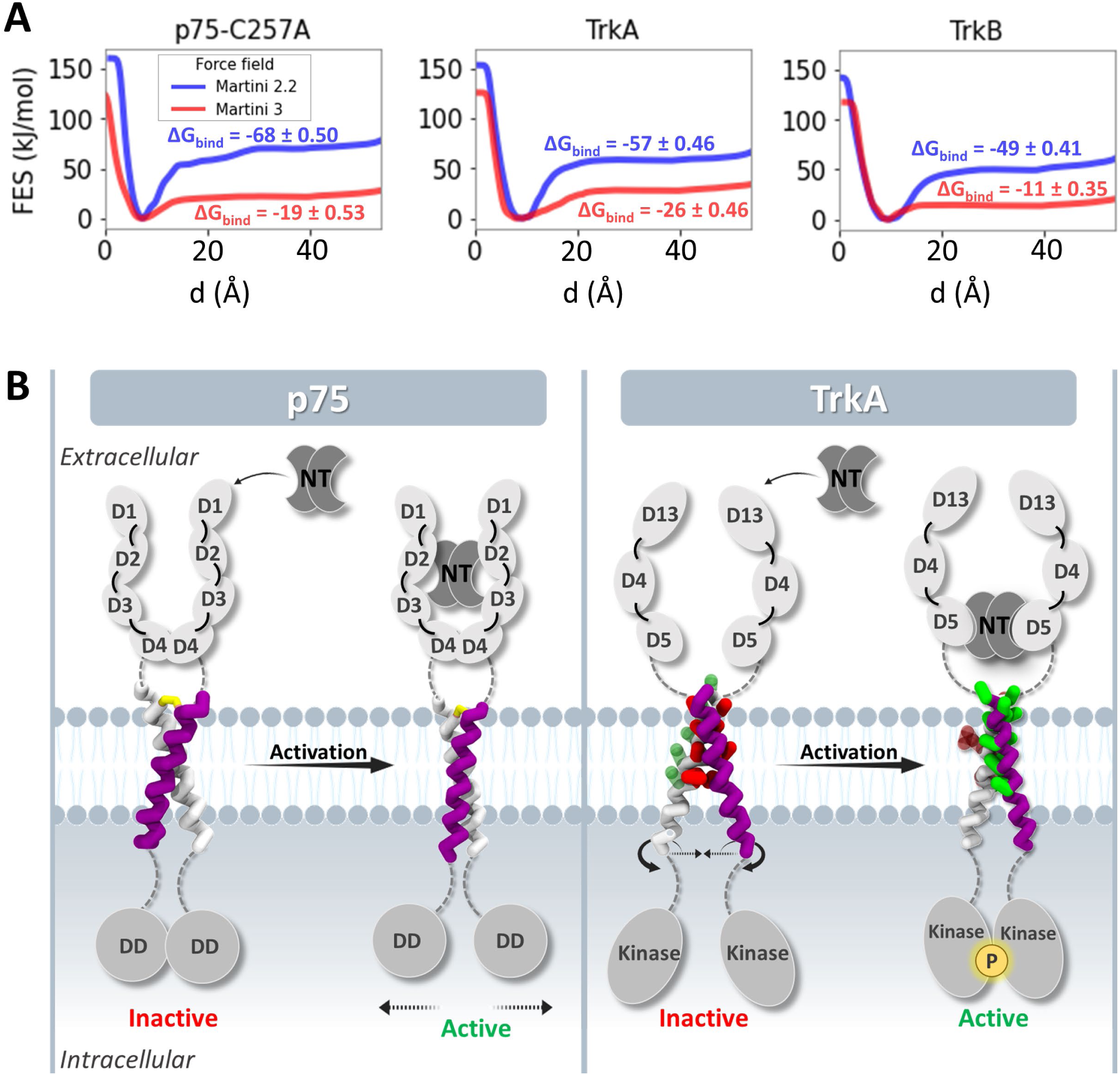
Simulations and computed free energy surfaces (FES) for helix association in a bilayer support different TM domain dimer-mediated activation mechanisms for the receptors. (A) Computed free energy surfaces (FES) for the association of the helices of the p75-C257A, TrkA and TrkB TM domain homodimers in a POPC bilayer show the lower stability of the TrkB TM domain dimer. 1D-FES as function of distance d between the helix centers of geometry (COGs) computed from the CG-MetaD simulations with Martini 2.2 and 3 (corresponding convergence plots are shown in Figures S5 and S6 and 2D-FES maps are shown in Figures S7 and S8). Corresponding values of ΔG_bind_ are given in kJ/mol. (B). The simulations indicate small fluctuations in the helix crossing angle, and consequently the distance between the helix C-termini, upon activation of p75, whereas the activation of TrkA involves a relative rotation of the helices (with little change in helix crossing angle) and a reduction in the distance between their C-termini. The different relative movements of the TM helices upon activation by NT binding lead to dissociation of the intracellular death domains (DD) of p75 and association followed by phosphorylation of the intracellular kinase domains of TrkA. The extracellular and intracellular domains and the neurotrophins (NTs) are shown schematically, while the transmembrane helices are shown in active and inactive arrangements derived from the molecular dynamics simulations (yellow: C257 cysteine, green: active and red: inactive interface sidechains).

In addition to the 1D FES over the biasing CV space, the FESs over all six geometric parameters that were used for the analysis of the unbiased CG simulations, were also calculated and plotted as 2D FESs (**Figures S7, S8**). Comparison of the FESs (**Figure S7)** and population densities (**Figure 3)** shows that similar helix arrangements were sampled for each system in the unbiased and MetaD simulations with Martini 3. The FES profiles differ between Martini 2.2 and 3 (**Figures S7, S8**) but, since the binding free energy calculated with Martini 3 is closer to the experimental values, the CG-MetaD simulations with Martini 3 are considered more realistic. The agreement between the CG-MetaD simulations and the unbiased simulations offers an additional validation of the sampling in both sets of simulations, and we conclude that the TM helix arrangements sampled in the unbiased simulations correspond to converged distributions.

## DISCUSSION

Transmembrane helix dimerization plays a crucial role in the activation and signal transduction of NT receptors and many other receptors, yet the molecular and mechanistic aspects of these processes remain poorly characterized. In this study, we investigated the TM domain dimerization of three NT receptors - p75, TrkA and TrkB – by employing a combination of unbiased all-atom (AA) and coarse-grained (CG) molecular dynamics (MD) simulations, along with enhanced-sampling CG-MetaD simulations. Not surprisingly, AA simulations failed to adequately sample the vast conformational space associated with TM helix dimerization whereas the CG simulations employing the Martini 3 force field gave a rather complete picture of the structural ensemble involved in TM helix dimerization. This observation is consistent with recent MD simulations of eleven other TM receptor dimers [35] that indicated broad applicability of CG simulations with the Martini 3 force field to sampling TM helix dimer arrangements. The results of our simulations underscore the critical influence of the lipid environment on the structural landscape of the TM domain dimers, revealing pronounced differences between micellar and bilayer environments. This observation for these NT receptors is supported by both experimental and computational studies that have highlighted the significance of the composition and structure of the lipid milieu for the transition between active and inactive states of bitopic membrane proteins.[36]–[38] Thus, considering that the phospholipid bilayers better represents the cell membrane than the micelles used in protein structure determination experiments and that the CG MD simulations enable broad sampling of the structural landscape of the TM domain dimers, we conclude that the CG MD simulations of the TM domain dimers in phospholipid bilayers should capture the native states of the receptors.

For the p75 low affinity NT receptor TM domain dimer, with its covalent disulfide bond at C257, a very narrow configurational space was sampled that was close to the NMR structure.[9] The helical dimer retained its right-handedness, with the only change during the simulation being in the crossing angle of the helices, which showed small fluctuations during the simulations. Interestingly, this motion is in agreement with FRET experiments, which have shown that p75 undergoes a conformational change upon NT binding in which C257 plays an important role.[18] The decrease in FRET upon NT binding suggested that the IC death domains dissociate when NT binds, and it has been proposed that NT binding induces a snail-tong-like movement of disulfide-linked p75, with the disulfide bond acting as the fulcrum, increasing the crossing angle upon activation.[18] However, these experiments did not provide any direct information on the motions of the TM domains whereas our simulations indicate small fluctuations of the crossing angle and, concomitantly, the distances between the termini of the helices to the active state of the receptor (**Figure 5**). The presence of the NT bound to the receptor could alter these fluctuations, leading to dissociation of the intracellular death domains. The complete change of the TM domain interface and the greater mobility in the simulations of the inactive C257A mutant show the key role of the disulfide bond in constraining the TM helix interactions.

Recent structural studies have proposed the existence of two distinct interfaces for the TrkA high-affinity NT receptor TM domain dimer: an active interface facilitating signal transduction and an inactive interface associated with quiescent states.[7] In the CG-MD simulations, both interfaces were sampled, shedding light on their respective structural dynamics and potential functional implications. Particularly intriguing is the potential impact of these interfaces on signal transduction pathways, where the conformational changes induced by the transition between the two states likely plays a crucial role in modulating downstream signaling events. From the simulation results, we suggest that a rotation of the TM helices and a close proximity of their C-termini accompany TrkA activation and could serve to facilitate autophosphorylation of the intracellular kinase domains, a pivotal step in the activation of TrkA-mediated signaling cascades (**Figure 5**). The structural ensemble of the active TrkA TM helical dimer derived from the simulations can potentially be used in structure-based drug design for the development of therapeutics that stabilize this state, as has been attempted for the TrkB receptor.[8], [21] Simulations with TrkA mutations at the active interface led to a change in the interacting interface, which provides an explanation for the drop in functional activity of these mutants observed experimentally.[7] The active interface observed in the simulations is also in agreement with disulfide cross-linking data for the TrkA V418C mutant.[7]

The activation dynamics of TrkB present differences with respect to TrkA. Only one active interface has been proposed for TrkB, in contrast to the dual interface model suggested for TrkA.[33] Furthermore, recent literature reveals discrepancies concerning the specific residues involved in this interface and whether the active TM domain dimer configuration is right- or left-handed.[8], [33] Additionally, the active interface of TrkB has been linked to an increase in the distance between C-terminal regions, in contrast to the activation mechanism that we propose for TrkA in which this distance decreases.[21], [39] In our simulations, we observed both right- and left-handed TrkB dimer configurations, with the latter being more prevalent. Notably, we initiated our simulations using a (left-handed) homology model derived from TrkA due to the absence of experimental data on the TrkB TM dimer structure when the simulations were performed. However, during the preparation of this manuscript, a newly released structural study proposed a right-handed NMR model of TrkB-TM dimer having an interface that overlaps with the most frequently observed arrangements in our simulations with residue S441 playing a key role in the interface.[33] Crucially, experimental evidence supports the notion that this interface corresponds to an active dimer configuration, thereby providing a validation of the ability of the CG MD simulations to sample different physiologically relevant states from the initial structural model. The interface between the TM helices of TrkB observed in our simulations is also similar to that of the TrkB-TM model bound to the antidepressant drug fluoxetine, which is a TrkB agonist.[8] In that study, fluoxetine binds to a helical arrangement where the A440xxxG444 motif is at the interface, which is also partially at the interface in our simulations, although our simulations show S441 rather than A440 at the interface. S441 is at the interface of the new NMR structure of the TrkB-TM dimer [33], and mutation of S441 to Ala or Ile leads to inhibition of TrkB signaling.[33] A further experimental validation of the arrangement of the TM domains in the CG-MD simulations corresponding to the active interface is that Y434, whose mutation to Cys leads to ligand-independent activation of TrkB,[40] is located at the most frequently observed interface in the simulations.

Even though a single dominant arrangement was observed for TrkB, the dimerization free energy of the helices was around half of that calculated for TrkA and p75-C257A, which suggests that other domains, such as the EC segments, or lipid components (like cholesterol) or small-molecules (like the agonist, fluoxetine) might be needed for additional stabilization of the active state of the dimer. The importance of the EC segments in TM domain regulation has already been shown for TrkA, for which the EC linker provides a coupling between the NT binding to the D5 domain and the TM helix rotation.[20] It should be noted that TrkB has a significantly longer EC linker than TrkA (51 residues for TrkB versus 35 residues for TrkA), and thus it would be of interest to understand how EC-TM coupling is achieved through this long disordered region.

A potential limitation of the work presented here is the simplicity of the phospholipid bilayer model. However, the simulations reveal arrangements of the TM domains consistent with experimental data and throw light on the different activation mechanisms of the three NT receptors studied. In further work, more complex membrane compositions including cholesterol, which has been proposed to have an important effect in TrkB-mediated signaling,[8] should be modeled. The complex composition of biological membranes, such as neuronal membranes, poses further demands on the computational sampling due to their heterogeneity and slowed dynamics and the current simulations provide important references for simulations with more complex bilayers. Moreover, including more domains of the receptors, ideally simulating the full-length glycoproteins, will be necessary to gain a complete understanding of the effects of the TM region (re)arrangements on signal propagation. Overall, though, the results of our simulations shed new light on the dynamic behavior of the TM domain homodimers of the p75, TrkA and TrkB NT receptors, reconcile the apparently contradictory experimental data, and point to how sequence differences can lead to distinct transmembrane signaling mechanisms. These differences provide an important basis for the engineering of protein signalling modules and for the design of peptide-based pharmaceuticals to specifically modulate NT signaling, an avenue that is being pursued for other TM helix receptors such as EGFR.[41], [42]

## MATERIALS AND METHODS

### Protein models

The NMR structures of the TrkA, p75 wild-type (WT) and C257A mutant homodimers of the TM domain with PDB IDs 2N90,[7] 2MIC[9] and 2MJO,[9] respectively, were retrieved from the RCSB.[43] Mutants containing single point mutations at six positions, V418A, V418C, G423I, L424A, A425I, F427A (underlined residues in **Figure 1B**), that were previously identified to belong to the active and inactive interfaces of the TrkA TM homodimer, [7] were modelled with Maestro (Schrödinger Suite [44]) in the NMR structure (PDB ID 2N90) of the TrkA TM homodimer.

For the TrkB TM homodimer, for which there was no experimentally determined structure available, bioinformatics web servers and databases were used to predict the TM and helical parts of the sequence. Specifically, UNIPROT,[45] ELM,[46] MEMSAT (PHYRE2),[47] MEMSAT-SVM,[48] TMpred,[49] TMHMM[50] and PredictProtein[51] were used for TM sequence prediction, while Jpred4,[52] PSIPRED,[53] NPSA-PRABI[54] and CFSSP[55] were used for secondary structure prediction (Table S2). From these predictions, the V433-R458 sequence was selected to be modeled as the TrkB TM helix. A homology model of the human TrkB TM homodimer was built with the AutoModel class of MODELLER v.9.23 [56] using the TrkA NMR structure (PDB ID 2N90) as a template. The TM sequences of TrkA, TrkB and p75 that were simulated are given in **Figure 1B**.

### Coarse-grained (CG) simulations

Coarse-grained (CG) molecular dynamics (MD) simulations of the TrkA, TrkB (1^st^ model), p75 WT and mutant TM dimers were performed in self-assembled DPC micelles, corresponding to the environment in which the NMR structures were determined, [7], [9] using the Martini 2.2 [22], [23] (Systems 3-12, **Table S1**) and Martini 3 [24] (Systems 13-16, **Table S1**) force fields. For the Martini 3 simulations, the protein-water Lennard-Jones potential well depths were scaled by a factor of 0.9, to allow self-assembly of the micelle around the TM protein, as we described recently.[31] The Martini 3 parameters for DPC from Ref. [31] were used. The initial atomistic models of the proteins were converted to Martini CG models with the martinize.py and martinize2.py scripts for Martini 2.2 and Martini 3, respectively. An elastic network model was used to preserve the secondary structure of the TM helices while retaining the intrinsic flexibility of the N- and C-terminal loop regions during the simulations. The elastic network applies additional harmonic restraints with an elastic force constant of 500 kJ/(mol·nm^2^) for Martini 2.2 and 700 kJ/(mol·nm^2^) for Martini 3, and a distance cutoff range of 5-9 Å. The CG homodimeric protein models were inserted in a simulation box together with 100 DPC molecules at random positions with a detergent-to-protein molar ratio (DPR) of 50:1, as used for the determination of the NMR structures of TrkA and of WT and mutant p75.[7], [9] The systems were solvated with the standard water model (NPW) with 10% of the anti-freeze water type (WF) for Martini 2.2 and the water model (W) for Martini 3, and neutralized by adding Na^+^ and Cl^−^ ions at 150 mM concentration. Additional test simulations (Systems 11-12, **Table S1**) were performed with an ionic strength of 24 mM and a detergent-to-protein molar ratio (DPR) of 20:1, corresponding to the experimental section of the 2N90 PDB file for the NMR structure, in order to examine the influence of these parameters on the sampled configurations of the TM homodimers.

All simulations were run with the GROMACS v.2020 MD engine.[57] Each simulation started with a steepest-descent energy minimization and was followed by a 2 μs equilibration in the NPT ensemble with the protein helical backbone restrained with a force constant of 4000 kJ/(mol·nm^2^), in order to retain the initial dimer arrangement, while the DPC micelles self-assembled around the protein TM domain. The self-assembly of the micelles was assessed by monitoring the radius of gyration of the DPC lipids, with constant low values indicating micelle formation (**Figure S9**). Additional test simulations (Systems 7-10, **Table S1**) were performed to examine the importance of the equilibrated micelle in the TM arrangements. Equilibration simulations were run at a constant temperature of 310 K, maintained using the velocity rescale thermostat [58] with a coupling constant of 1 ps and at a constant pressure of 1 bar, maintained with the Berendsen barostat.[59] A coupling constant of 4 ps was used to maintain isotropic pressure coupling with a compressibility of 4.5·10^-5^. A time step of 20 fs was applied. The non-bonded interactions were treated with a reaction field for Coulomb interactions, and the cutoff distance for these and for van der Waals interactions was 1.1 nm. Twenty replica equilibration simulations were run for each of the four systems (TrkA, TrkB, WT and mutant p75) starting from different initial protein and lipid positions, as well as different initial velocities, and ten replicas in which equilibrated micelles had been successfully formed by the end of the 2 μs equilibration were progressed to production simulations. These were run for fully unrestrained systems for a duration of 20 μs. Production simulations were run in the NPT ensemble at a constant temperature of 310 K, maintained using the velocity rescale thermostat with coupling constant of 1 ps and at a constant pressure of 1 bar, maintained with the Parrinello-Rahman barostat.[60], [61] A coupling constant of 12 ps was used to maintain isotropic pressure coupling with a compressibility of 3·10^-4^. A time step of 20 fs was applied. The non-bonded interactions were treated with a reaction field for Coulomb interactions, and the cutoff distance for these and for van der Waals interactions was 1.2 nm.

Additional CG simulations of the TrkA WT and V418A, V418C, G423I, L424A, A425I and F427A mutants, TrkB WT, p75 WT and C257A mutant TM dimers were performed in POPC bilayers with Martini 3 (Systems 17-26, **Table S1**). The protein models were embedded in an 80 Å x 80 Å POPC bilayer grid using the insane (INSert membrANE) script. [62] The systems were energy minimized with the steepest-descent algorithm and ten replica simulations starting from different initial velocities were equilibrated with the protein restrained as described above and then the restraints were removed in the subsequent production simulations. All the simulation parameters were the same as for the simulations with the Martini 2.2 force field in DPC micelles with the exception that the cutoff distance for Coulomb and van der Waals interactions was 1.1 nm for the production runs and semi-isotropic pressure coupling was used. For each of these systems, 10 replicas were run for 20 μs each. Overall, a total of 4.2 ms of unbiased CG MD simulations were performed for the different systems and lipid environments.

### Atomistic simulations

All-atom (AA) MD simulations of the TrkA and TrkB TM dimers in DPC micelles were performed using the CHARMM36m force field [30] and the CHARMM-modified TIP3P water model (Systems 1-2, **Table S1**). The simulations in DPC micelles were initiated from the ten replica CG systems containing self-assembled, equilibrated micelles, after back-mapping to atomistic detail using the backward.py script developed by the Martini team. [63] For this purpose, a mapping was constructed for DPC lipids (**Supplementary Data 1**) to define the CG beads that correspond to specific atoms. All simulations were run with the GROMACS v.2020 MD engine. Each simulation started with a steepest-descent energy minimization and was followed by one NVT and three NPT equilibrations in which gradual removal of restraints on protein atoms took place. Specifically, the first NVT equilibration was run at 310 K for 200 ps with the protein helical backbone (BB) harmonically restrained with a force constant of 4000 kJ/(mol·nm^2^) and the side chains (SC) restrained with a force constant of 2000 kJ/(mol·nm^2^). The subsequent three (A, B and C) NPT equilibrations were run at 310 K and 1 bar pressure for 1 ns each with diminishing BB beads and SC restraints with force constants of A: BB:4000 kJ/(mol·nm^2^), SC:2000 kJ/(mol·nm^2^), B: BB:2000 kJ/(mol·nm^2^), SC:1000 kJ/(mol·nm^2^), and C: BB:1000 kJ/(mol·nm^2^), SC:500 kJ/(mol·nm^2^). For the first three equilibrations, the Berendsen thermostat and barostat were used and for the last equilibration, the Nosé-Hoover thermostat [64], [65] and the Parrinello-Rahman barostat were used. A coupling constant of 5 ps was used to maintain isotropic pressure coupling with a compressibility of 4.5·10^-5^. A time step of 1 fs was applied in the first equilibration step and then the time step was increased to 2 fs for the next steps. The non-bonded electrostatic interactions were treated with PME with a cutoff distance at 1.2 nm. Non-bonded van der Waals interactions were calculated with a cutoff of 1.2 nm and a switching distance of 1.0 nm. The LINCS algorithm [66] was employed to constrain the length of all hydrogen-containing bonds. Production runs were subsequently run for 1 μs for each of the 10 replicas with the protein completely unrestrained. The same simulation parameters were used as for the last equilibration step. A total of 20 μs of AA simulations were performed.

### Coarse-grained metadynamics (CG-MetaD) simulations

Well-tempered metadynamics [67] simulations of the TrkA, TrkB and mutant p75-C257A TM dimers in a POPC bilayer were performed at the CG level with the Martini 2.2 and Martini 3 force fields (Systems 27-32, **Table S1**) to calculate the free energy surfaces for dimerization. The simulations were run with GROMACS v.2020 patched with Plumed v.2.7.[68] The protein models were embedded in an 80 Å x 80 Å POPC bilayer grid using the insane script.[62] Energy minimization was performed with the steepest-descent algorithm and then equilibration with the protein restrained as described above in the “Coarse-grained (CG) simulations” section. Then, the CG-MetaD production simulations were run after removing the restraints.

The distance (d) between the COGs of the backbone (BB) beads of the two helix monomers was used as a biasing collective variable (CV) (**Figure 2B**). A Gaussian potential was deposited on the CV space every 5000 steps with decreasing height W = W_0_e^-V(s,t)/(f-1)T^, where W_0_ = 0.05 kJ/mol is the initial Gaussian height, T = 310 K is the simulation temperature, f = 10 is the bias factor, and V(d,t) is the bias potential as a function of the CV(s) and the time. The Gaussian width was set to 0.05 nm, which was approximately half of the CV fluctuations in the unbiased simulations. These metadynamics parameters have been previously used successfully for the reconstruction of the free energy surface of the EGFR TM domain dimerization.[25] The rest of the simulation parameters were the same as for the unbiased CG simulations of the membrane systems described above. The CG-MetaD simulations were run for more than 100 μs for each system, resulting in a total of ∼760 μs for all the CG-MetaD simulations.

### Analysis of simulations

Trajectory frames were saved every 1 ns. All frames were post-processed with the GROMACS gmx trjconv tool to remove periodic boundary condition (pbc) jumps, center the proteins in the simulation box and ensure that molecules are not fragmented. The systems were aligned to the backbone of the first TM domain helix throughout the trajectories. Graphical rendering of protein structures from the trajectories was performed with VMD 1.9.3. [69]

#### Geometric parameters

Parameters were calculated that describe the various arrangements of the TM helices that were sampled during the simulations. These parameters were: (1) the distance between the centers of geometry (COG) of the backbone (BB) beads of the two helices, (2) the crossing angle between the two helices defined as the dihedral angle formed by the COGs of the BB beads of the two helices and the COGs of the BB beads of the N-terminal halves of the two helices, (3) the phase and position, which correspond to the rotation around each helix, defined as the dihedral angle formed by the upper COG of the backbone (BB) beads of one helix, the COG of the BB beads of the same helix, the COG of the backbone (BB) beads of the second helix and one residue, which was A428 for TrkA, G444 for TrkB, and V265 for p75 and p75-C257A, (4) the distances between the N-termini and between the C-termini of the two helices, defined by the COGs of the BB beads of the first and last residues (Figure 2B and Figure 3). For the calculation of these parameters, MDAnalysis [70], [71] was used with the numpy.linalg.norm function of the Numpy library.[72] These parameters were calculated for all frames and all replicas.

#### Kernel Density Estimation (KDE)

The population density distributions of the geometric parameters were visualized with Kernel Density Estimation (KDE), which smooths the density by summing kernels on the sampled data points. The 2D population density distributions of two parameters were calculated and visualized with the seaborn.kdeplot function of Seaborn using the default bandwidth value of 1.0.[73]

#### Contact analysis

The residues from each helix that form contacts with the residues of the other helix were calculated with MDAnalysis. A contact was defined as formed when any CG bead of a residue was within a distance of less than 6 Å of another bead. The contacts were calculated for all frames of all replicas. The data were post-processed with the Pandas library [74], [75] and the results were visualized with Matplotlib.[76]

#### CG-MetaD analysis

To assess the convergence of the CG-MetaD simulations, several quantities were monitored: (1) the time evolution of the distance between the COGs of the BB beads of the two helices was calculated with Plumed to assess diffusion in the biasing CV, (2) the decrease of the height of the deposited Gaussian potentials over time, (3) the time evolution of the free energy difference between the bound and unbound states in the biasing CV space, (4) the average free energy difference across blocks and the error as a function of block size using the block-analysis technique and calculating the error every 10 blocks from one to 1000 blocks.

For the 2D energy landscapes, a reweighting protocol was used to remove the bias and compute the Boltzmann distribution along the same geometric parameters as were monitored in the unbiased simulations. Thus, the free energy surface (FES) was reconstructed as a function of both the original biasing CV (the interhelical distance d) and the interhelical crossing angle (Ω). Additionally, the FES was reconstructed as a function of the other descriptors for helix dimer arrangement described in the “Geometric parameters” section. For the FES, these parameters, their histograms, and the corresponding conversion to free energy were calculated with Plumed.

## Supplementary Materials

Figures S1-S9

Tables S1-S2

Supplementary Data 1: Mapping between Martini coarse-grained beads and atoms for a DPC lipid molecule.

Supplementary References

## Acknowledgments

The authors acknowledge provision of computing resources by the state of Baden-Württemberg through bwHPC and the German Research Foundation (DFG) through grant INST 35/1134-1 FUGG. We thank Stefan Richter for technical support and Mislav Brajkovic, Jonathan Teuffel, Giulia Paiardi, Hendrik Jung and Balazs Fabian for comments on the manuscript.

## Author contributions

Examples:

Conceptualization: CA, RCW

Methodology: CA, ACC, AT, RCW

Investigation: CA, ACC, AT

Visualization: CA, ACC, AT

Funding acquisition: RCW

Project administration: RCW

Supervision: RCW

Writing – original draft: CA, ACC, AT, RCW

Writing – review & editing: CA, ACC, AT, RCW

## Data and materials availability

Input files, the structures shown in figures and representative trajectories are available on Zenodo at doi:10.5281/zenodo.12192514. All other data are available from the authors upon request.

