## Supplementary Material for "Simulation of neurotrophin receptor transmembrane helix interactions reveals active states and distinct signaling mechanisms"

### Supplementary Materials

**Authors:** Christina Athanasiou[1,2,3,†], Ainara Claveras Cabezudo[1,2,†], Alexandros Tsengenes[1,2,3], Rebecca C. Wade[1,2,4,5,*]

**Affiliations:**

[1] Molecular and Cellular Modeling Group, Heidelberg Institute for Theoretical Studies (HITS); Heidelberg, Germany.

[2] Faculty of Biosciences, Heidelberg University; Heidelberg, Germany.

[3] Heidelberg Biosciences International Graduate School, Heidelberg University; Heidelberg, Germany.

[4] Center for Molecular Biology of Heidelberg University (ZMBH), DKFZ-ZMBH Alliance, Heidelberg, Germany.

[5] Interdisciplinary Center for Scientific Computing (IWR), Heidelberg University; Heidelberg, Germany.

†These authors contributed equally to this work

### Contents

Figures S1-S9
Tables S1-S2
Supplementary Data file 1: Mapping between Martini coarse-grained beads and atoms for a DPC lipid molecule.
Supplementary References

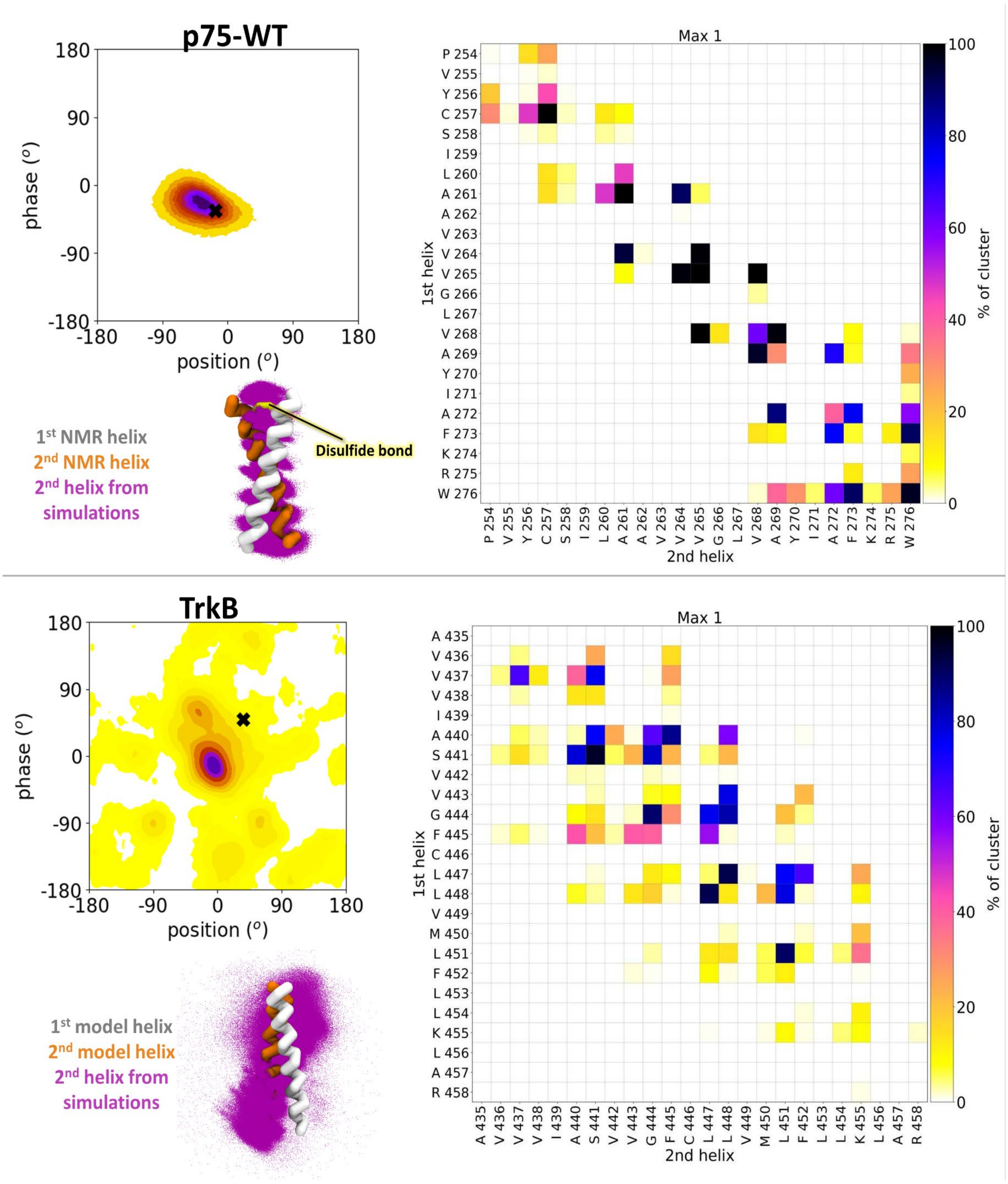


**Figure S1: Helix contact maps for the dominant arrangements of the p75 and TrkB TM homodimers in a POPC bilayer environment.** In each case, the heatmaps on the left show the population density from the CG simulations with the Martini 3 force field for p75 and TrkB in POPC bilayers projected onto the 2D phase-position space (the reference residues used for the phase-position calculations are V265 for p75 and G444 for TrkB and the NMR structure for p75-WT and the homology model of TrkB, from which the simulations were started, are indicated with black crosses (**X**); these heatmaps are also shown in Fig 3E and 3H, respectively). The helix contact maps that correspond to the dominant maximum in the phase-position heatmap are shown on the right. A contact is defined when two CG beads are within 6 Å distance. The contact occupancy within the cluster of arrangements that corresponds to the maximum is shown for each residue pair. The corresponding TM domain arrangements are shown with all the frames aligned to the 1st helix (white). The 2nd helix from the NMR structure (p75) or initial homology model (TrkB) is shown in orange for reference, and the 2nd helix from the simulation frames is shown in purple dots.

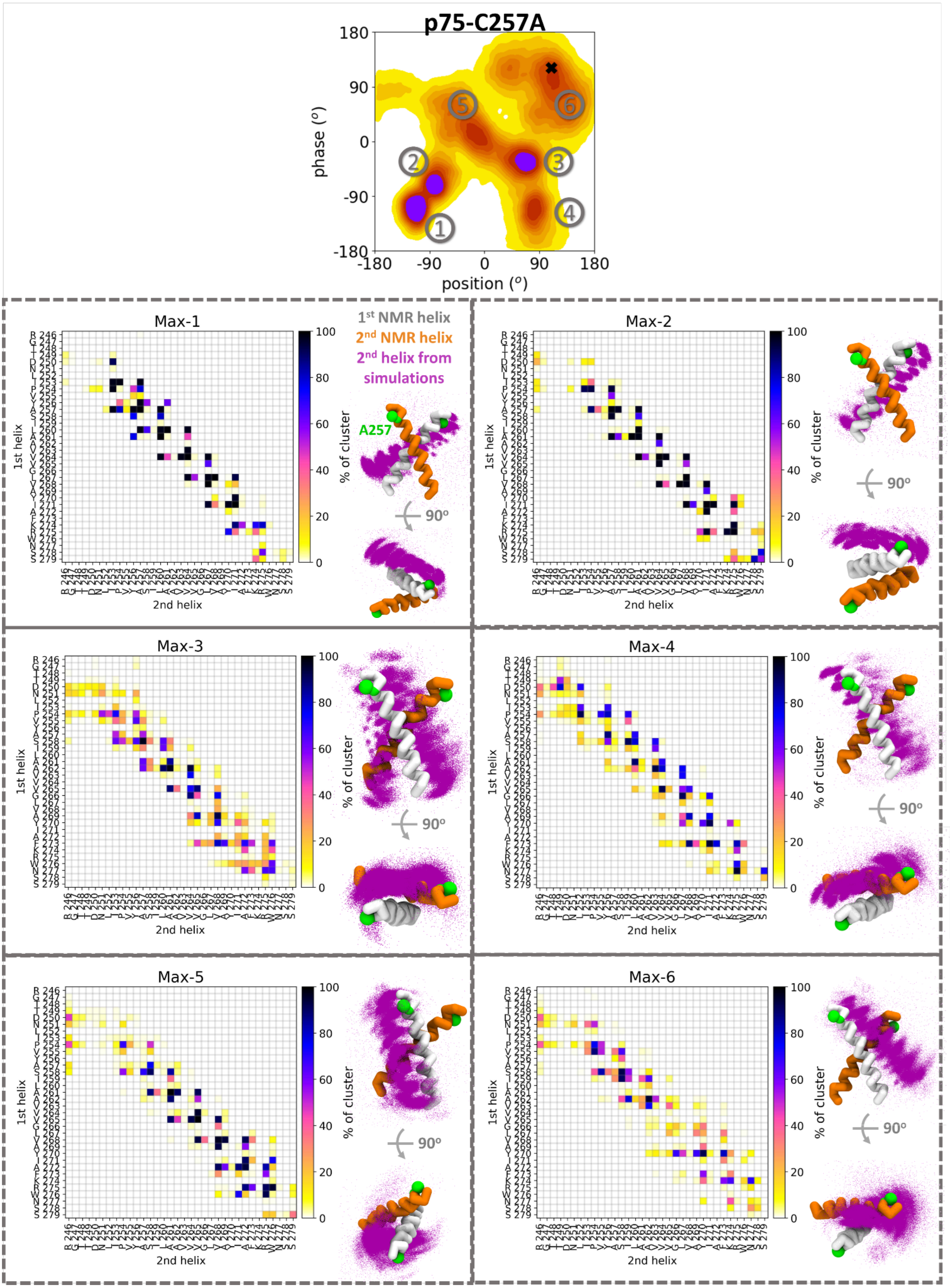
p75-C257A
phase (°)
position (°)
Max-1
Max-2
Max-3
Max-4
Max-5
Max-6
1st NMR helix
2nd NMR helix
2nd helix from simulations
A257
1st helix
2nd helix
% of cluster
90°

**Figure S2: Helix contact maps of dominant arrangements of the p75-C257A TM homodimer in a POPC bilayer environment.** The upper plot shows the population density heatmap from the CG simulations with the Martini 3 force field for p75-C257A in a POPC bilayer projected onto the 2D phase-position space (the reference residue used for the phase-position calculations was V265 for p75-C257A, the NMR structure is indicated with a black cross (**X**); this heatmap is also shown in Fig 3F). The maxima observed in the heatmap are numbered and analyzed in the lower plots. The helix contact maps that correspond to each of the dominant maxima in the phase-position heatmap are shown together with a visualization of the TM domain arrangements. A contact is defined when two CG beads are within 6 Å distance. The contact occupancy within the cluster of arrangements that correspond to the given maximum is shown for each residue pair. The corresponding TM domain arrangements are shown with all the frames aligned to the 1st helix (white). The 2nd helix from the NMR structure is shown in orange for reference, and the 2nd helix from the simulation frames in the cluster of arrangements that correspond to the respective maximum is shown by purple dots.

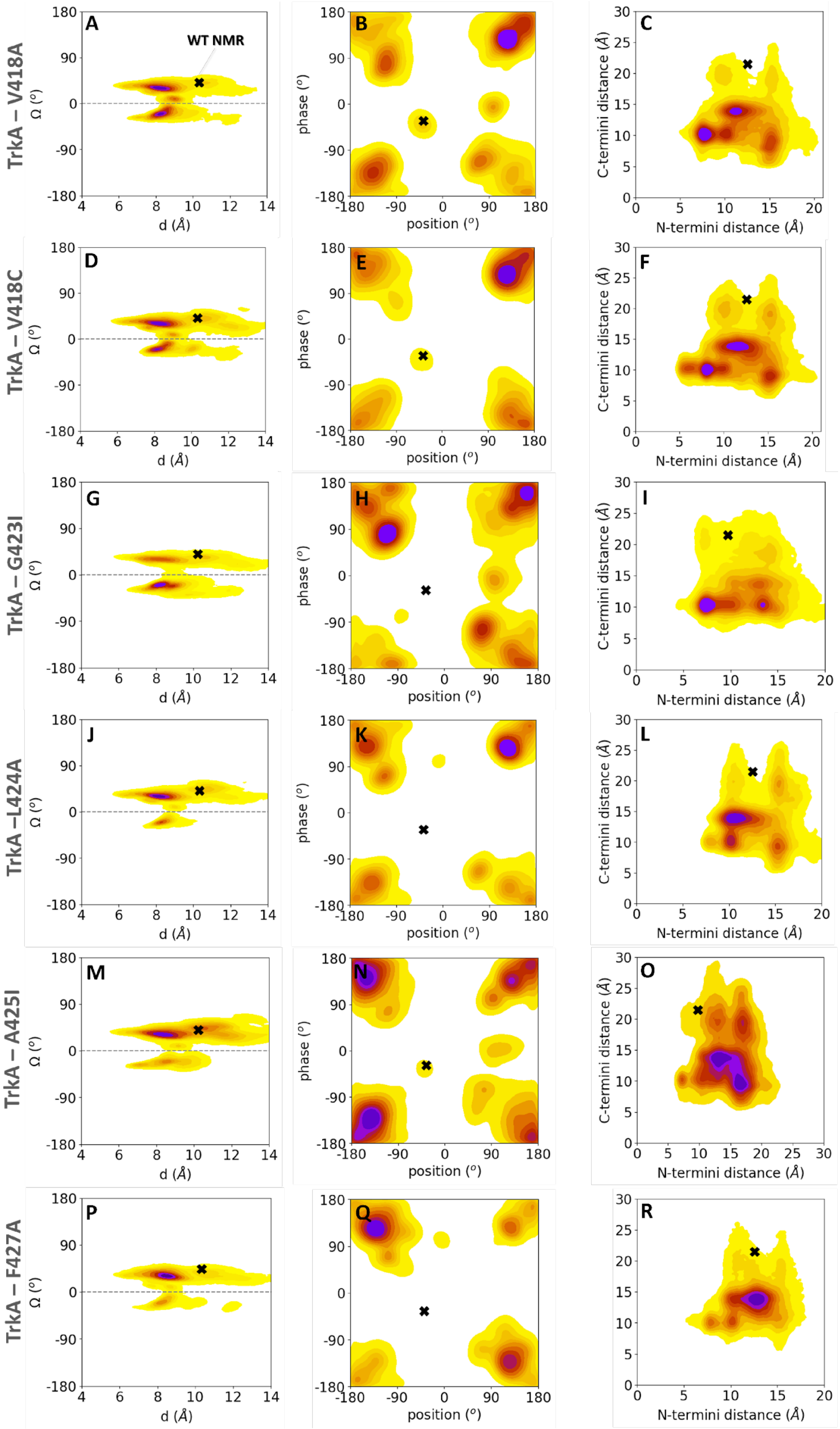
TrkA – V418A
TrkA – V418C
TrkA – G423I
TrkA –L424A
TrkA – A425I
TrkA – F427A
A
B
C
D
E
F
G
H
I
J
K
L
M
N
O
P
Q
R
WT NMR
Ω (°)
d (Å)
phase (°)
position (°)
C-termini distance (Å)
N-termini distance (Å)

**Figure S3: TM helix arrangements in a POPC bilayer environment of six TrkA mutants in the space of six geometric parameters.** Population density heatmaps from CG simulations with the Martini 3 force field of the TrkA V418A, V418C, G423I, L424A, A425I, F427A mutants in a POPC bilayer are shown projected onto the 2D spaces for d-Ω, phase-position (the reference residue used for the phase-position calculations is A428), and C-termini and N-termini distances. The population maxima are denoted in dark purple and the color fades to yellow for states of lower probability. The NMR structure for WT TrkA is indicated with a black cross (**X**). Positive Ω values indicate left-handed helical dimers (L), while negative values correspond to right-handed dimers (R). See Fig. 5 for further analysis of the TrkA V418A and VF427A mutants.

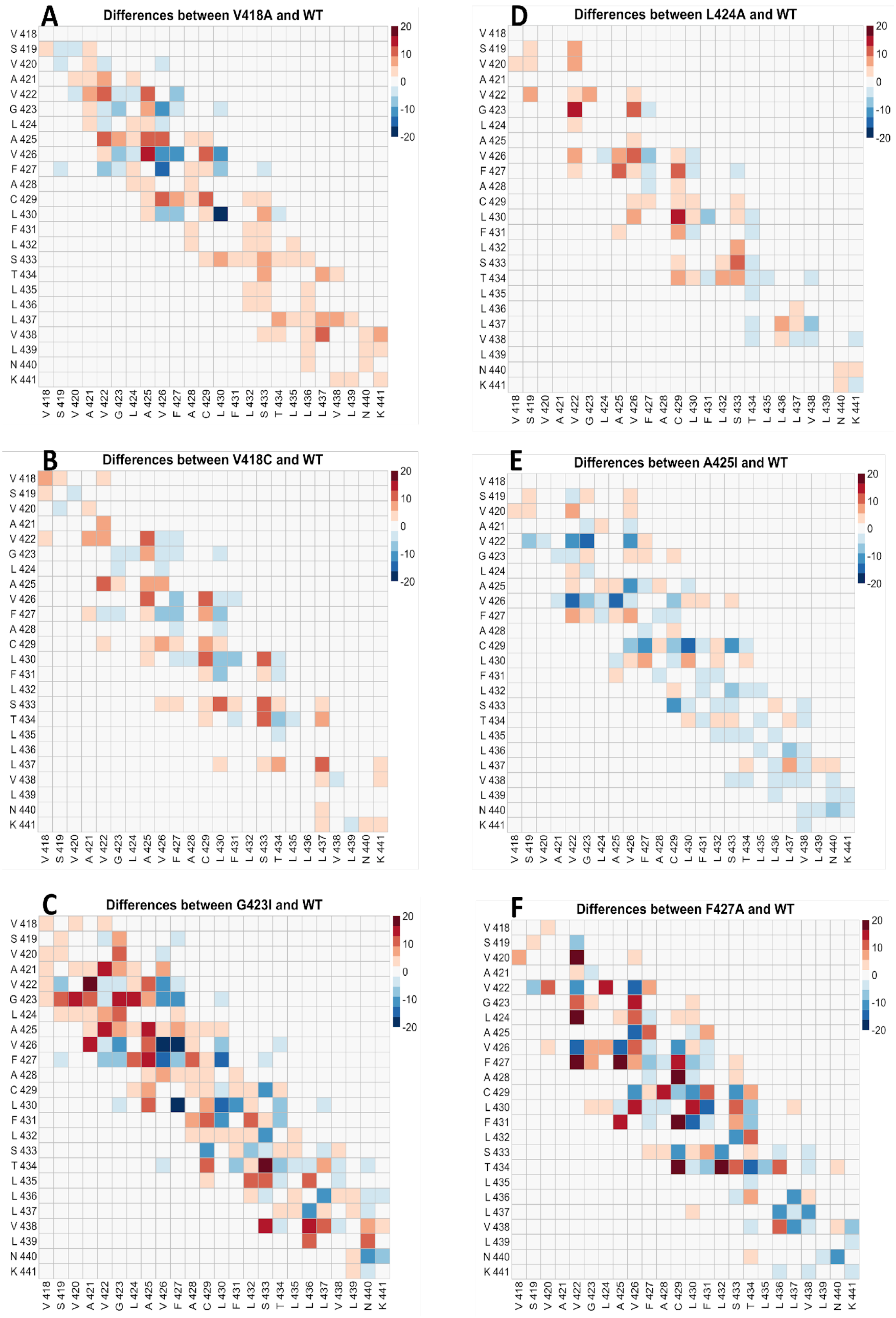


**Figure S4: Differences in TM helix contact maps for six TrkA mutants compared to WT TrkA.** The contacts of the WT TrkA were subtracted from the contacts of the TM helices of the TrkA V418A, V418C, G423I, L424A, A425I and F427A mutants (blue: fewer contacts in the mutant, red: more contacts in the mutant compared to WT). See Fig. 5 for further analysis of the TrkA V418A mutant.

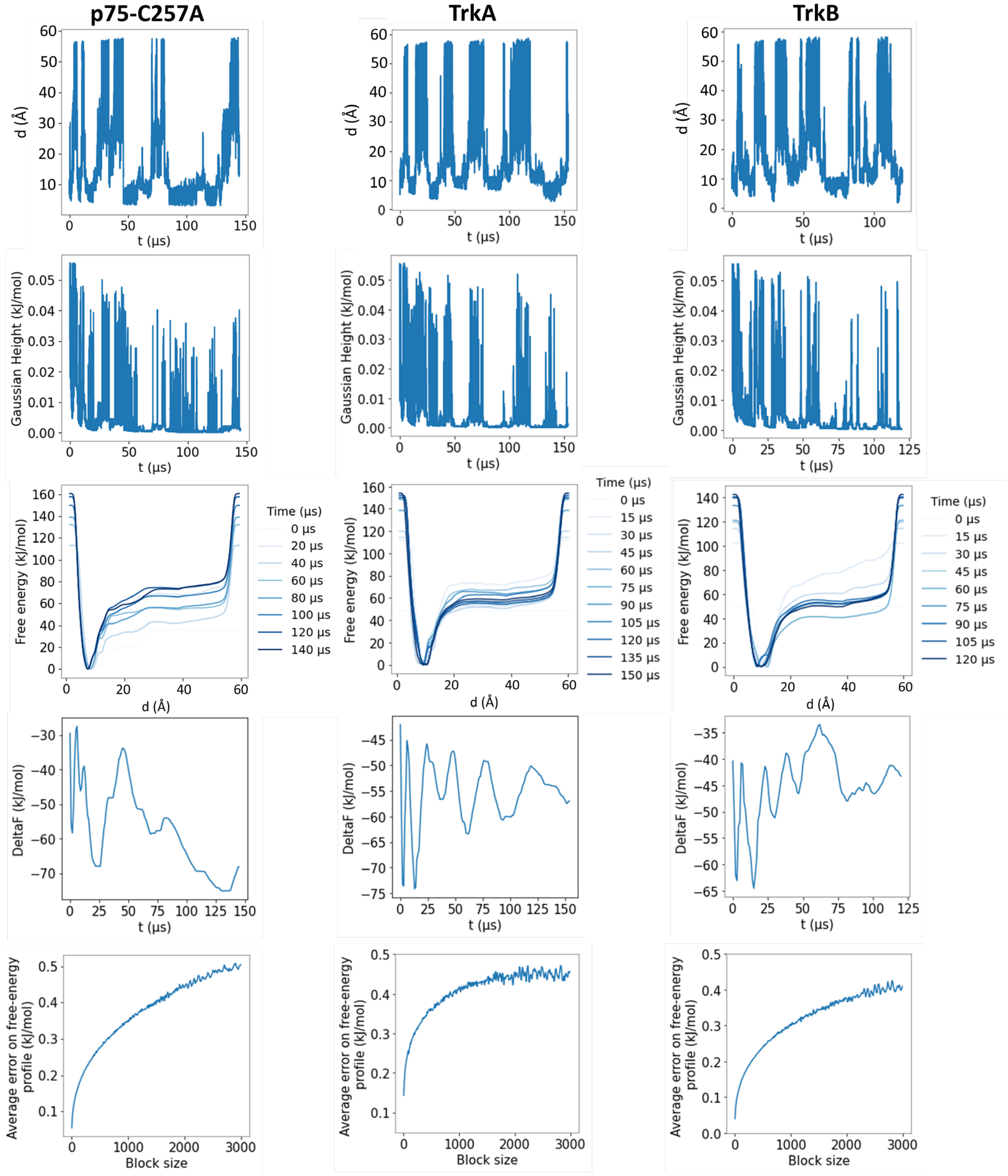


**Figure S5: Evaluation of the convergence of the CG-MetaD simulations with the Martini 2.2 force field for the p75-C257A, TrkA and TrkB TM helix dimer systems.** The properties monitored as a function of simulation time are from top to bottom: interhelical distance, d, (the biasing CV), the Gaussian height, the free energy along CV space, the free energy difference between the bound state (d ≤ 30 Å) and the unbound state (30 < d ≤ 40 Å), and the average error in the free energy, shown as a function of simulation block size. The diffusive behavior of the sampling in the CV indicates convergence. The free energy surfaces are shown in Figure 6.

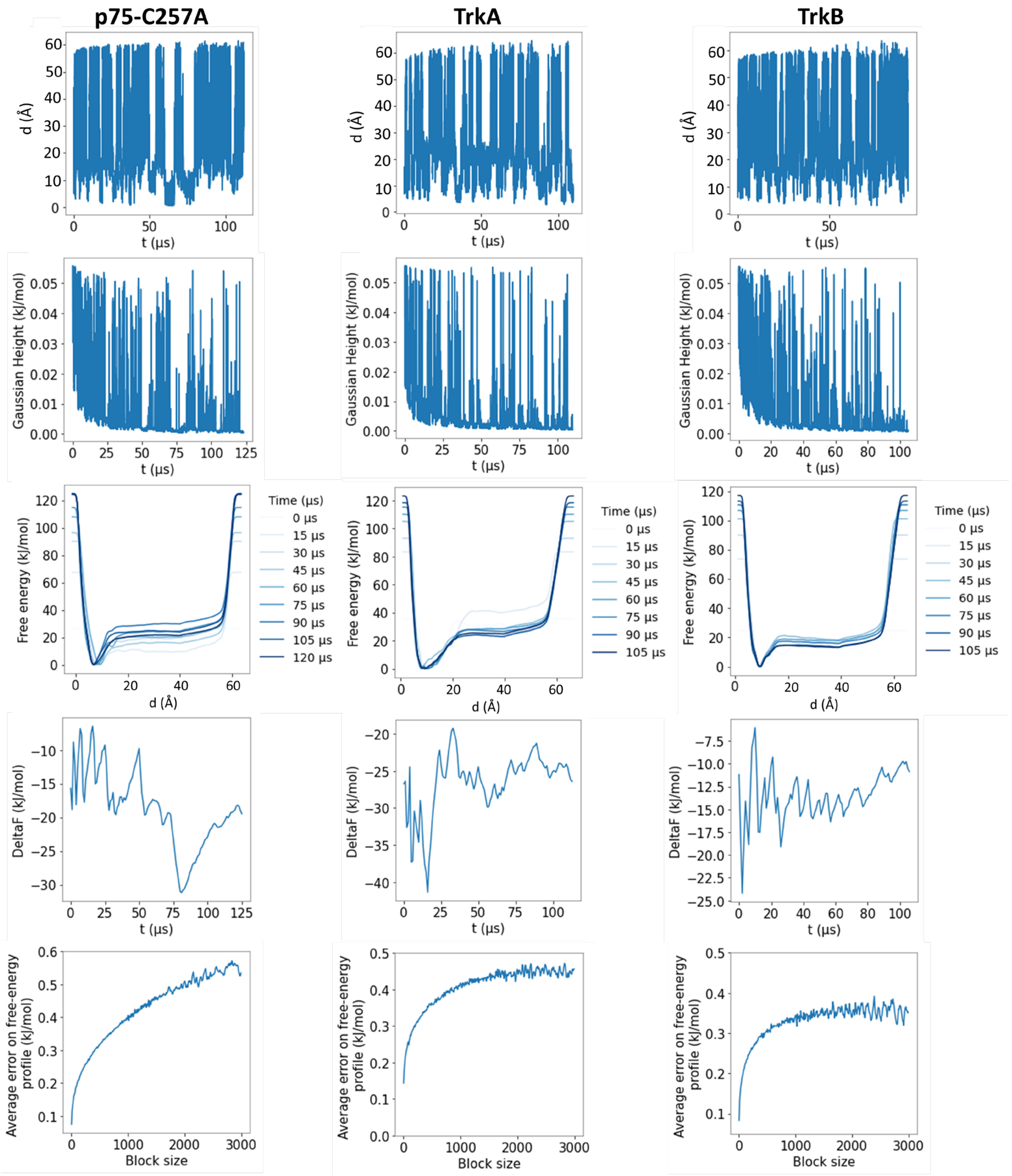


**Figure S6: Evaluation of the convergence of the CG-MetaD simulations with the Martini 3 force field for the p75-C257A, TrkA and TrkB TM helix dimer systems.** The properties monitored as a function of simulation time are from top to bottom: interhelical distance, d, (the biasing CV) , the Gaussian height, the free energy along CV space, the free energy difference between the bound state (d ≤ 30 Å) and the unbound state (30 < d ≤ 40 Å), and the average error in the free energy, shown as a function of simulation block size. The simulations showed a diffusive exploration of the CV space with Gaussian heights decreasing over time. The diffusion over CV space was higher for the Martini 3 simulations than the Martini 2 simulations, indicating better convergence. The free energy surfaces are shown in Figure 6.

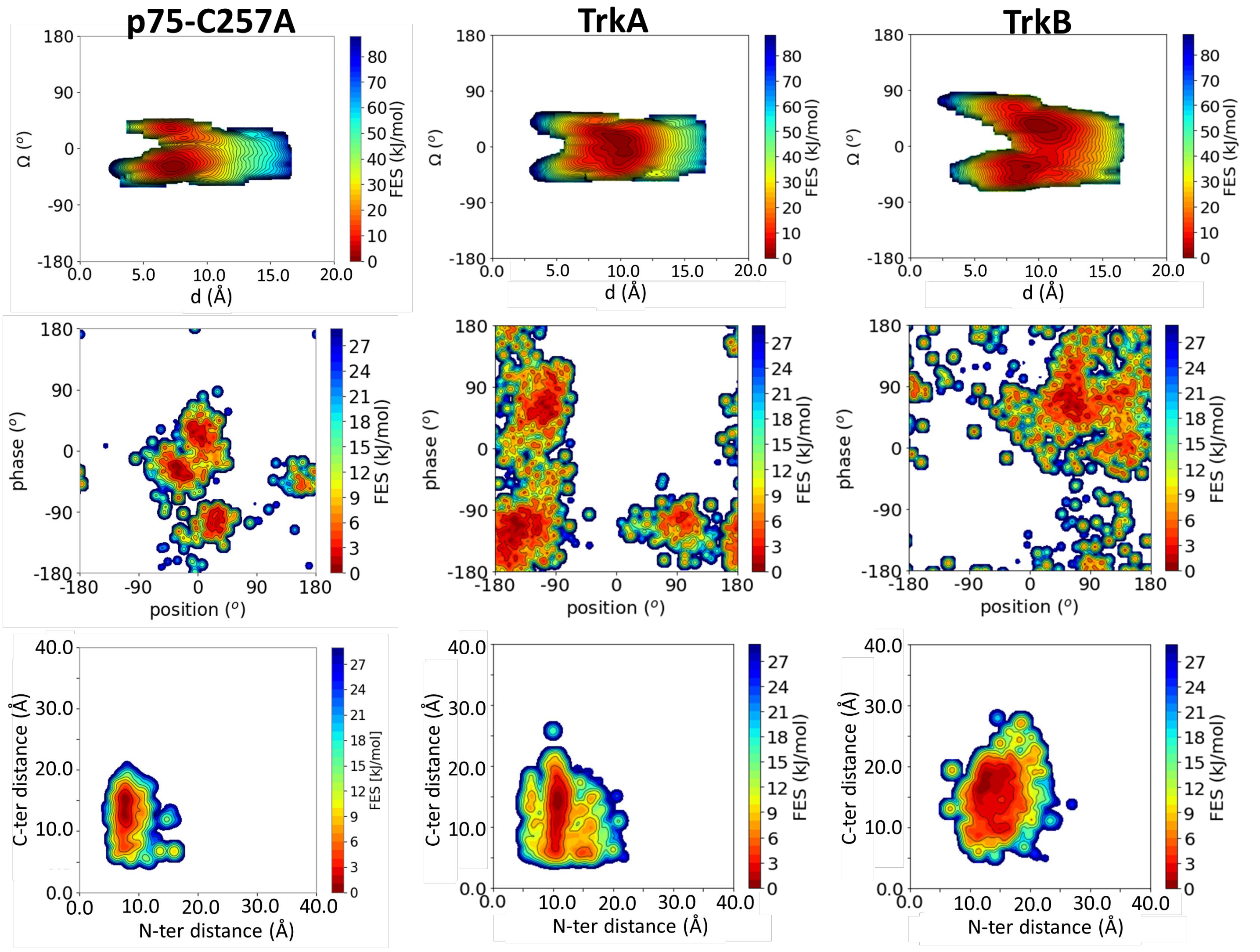


**Figure S7: Free energy surfaces (FES) from the CG-MetaD simulations with Martini 2.2 for the P75-C257A, TrkA and TrkB TM helix dimer systems.** The FES was calculated by reweighting in the space of crossing angle (Ω) – interhelical distance (d), phase-position and N- and C-termini distances. Isosurfaces are plotted every 5 kJ/mol for Ω-d, and every 3 kJ/mol for phase-position and C- and N-termini distances.

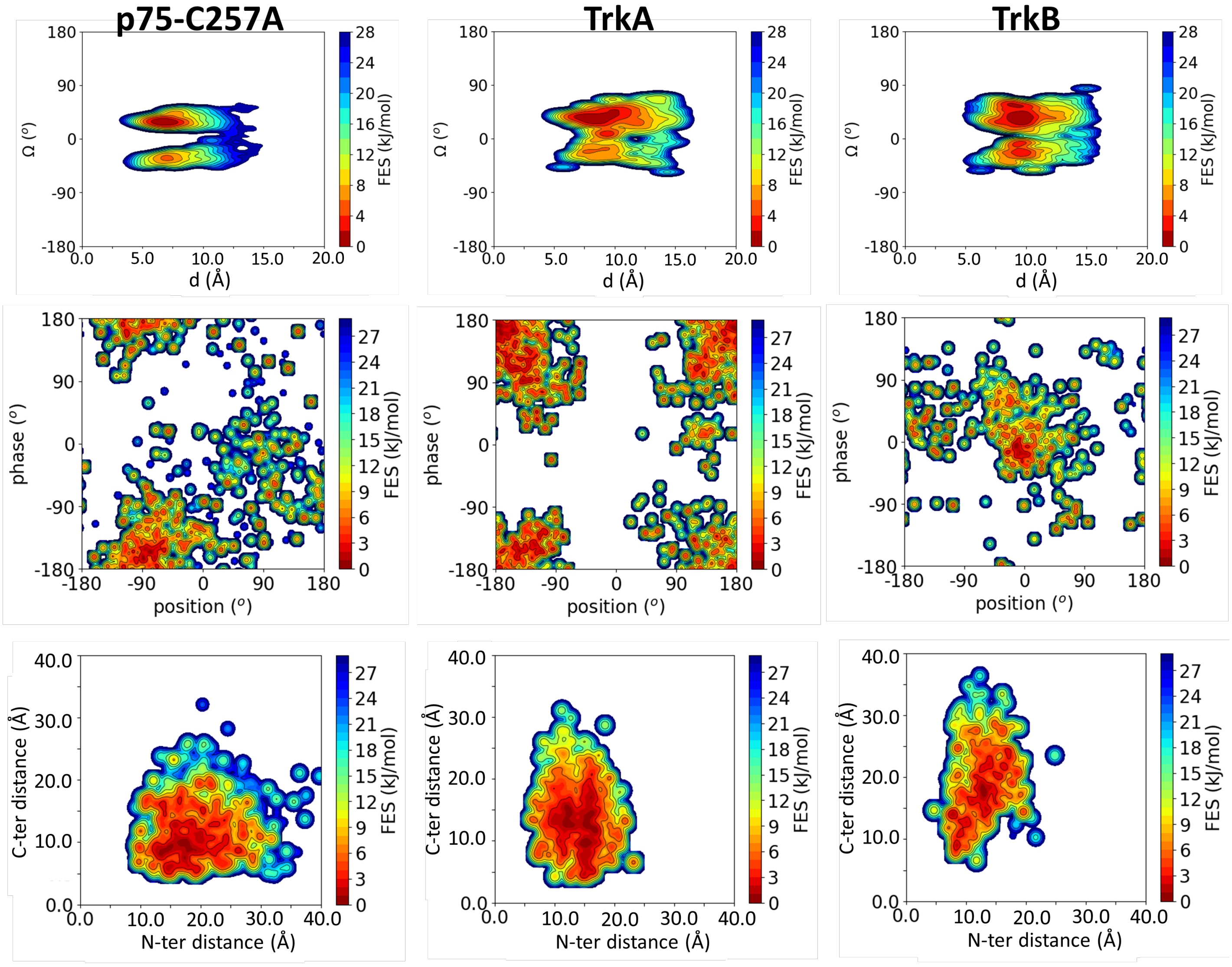


**Figure S8: Free energy surfaces (FES) from the CG-MetaD simulations with Martini 3 for the P75-C257A, TrkA and TrkB TM helix dimer systems.** The FES was calculated by reweighting in the space of crossing angle (Ω) – interhelical distance (d), phase-position and N- and C-termini distances. Isosurfaces are plotted every 5 kJ/mol for Ω-d, and every 3 kJ/mol for phase-position and C- and N-termini distances. The corresponding 1-D free energy surfaces are shown in Figure 6.

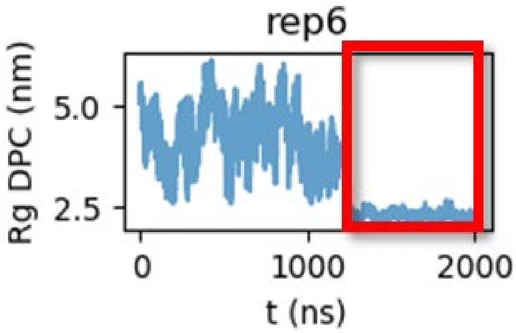


**Figure S9: Example of monitoring of the radius of gyration (Rg) of the DPC lipids during the equilibration of a micelle self-assembly replica CG simulation of a Trk TM helix dimer system**. Low and constant values of Rg indicate formation of a micelle. The red box indicates the part of the equilibration simulation where a micelle has been formed. All 20 equilibration simulation trajectories for each of the TrkA and TrkB TM helix dimer systems were also examined visually for micelle formation.

**Table S1: Overview of all simulations performed.** Test simulations are denoted with *.[1] Dimers of the transmembrane (TM) domain were simulated with DPC or POPC lipids. For systems 11 and 12, I is the ionic strength and DPR is the detergent-to-protein ratio. The simulation types are all-atom (AA), coarse-grained (CG) with Martini 2.2 (M2) or Martini 3 (M3), and coarse-grained metadynamics (CG-MetaD).

| System No | Protein system | Lipid environment | Simulation type | Replicas | Time/Replica (μs) |
|---|---|---|---|---|---|
| 1 | TrkA TM | DPC micelle | AA | 10 | 1 |
| 2 | TrkB TM | DPC micelle | AA | 10 | 1 |
| 3 | TrkA TM | DPC micelle | CG-M2 | 10 | 20 |
| 4 | TrkB TM | DPC micelle | CG-M2 | 10 | 20 |
| 5 | p75-WT TM | DPC micelle | CG-M2 | 10 | 20 |
| 6 | p75-C257A TM | DPC micelle | CG-M2 | 10 | 20 |
| 7* | TrkA TM | DPC non-equil. micelle | CG-M2 | 10 | 5 |
| 8* | TrkB TM | DPC non-equil. micelle | CG-M2 | 10 | 5 |
| 9* | p75-WT TM | DPC non-equil. micelle | CG-M2 | 10 | 5 |
| 10* | p75-C257A TM | DPC non-equil. micelle | CG-M2 | 10 | 5 |
| 11* | TrkA TM | DPC micelle, I = 24 mM | CG-M2 | 10 | 20 |
| 12* | TrkA TM | DPC micelle, I = 24 mM, DPR = 20:1 | CG-M2 | 10 | 20 |
| 13 | TrkA TM | DPC micelle | CG-M3 | 10 | 20 |
| 14 | TrkB TM | DPC micelle | CG-M3 | 10 | 20 |
| 15 | p75-WT TM | DPC micelle | CG-M3 | 10 | 20 |
| 16 | p75-C257A TM | DPC micelle | CG-M3 | 10 | 20 |
| 17 | TrkA TM | POPC bilayer | CG-M3 | 10 | 20 |
| 18 | TrkB TM | POPC bilayer | CG-M3 | 10 | 20 |
| 19 | p75-WT TM | POPC bilayer | CG-M3 | 10 | 20 |
| 20 | p75-C257A TM | POPC bilayer | CG-M3 | 10 | 20 |
| 21 | TrkA-V418A TM | POPC bilayer | CG-M3 | 10 | 20 |
| 22 | TrkA-V418C TM | POPC bilayer | CG-M3 | 10 | 20 |
| 23 | TrkA-G423I TM | POPC bilayer | CG-M3 | 10 | 20 |
| 24 | TrkA-L424A TM | POPC bilayer | CG-M3 | 10 | 20 |
| 25 | TrkA-A425I TM | POPC bilayer | CG-M3 | 10 | 20 |
| 26 | TrkA-F427A TM | POPC bilayer | CG-M3 | 10 | 20 |
| 27 | TrkA TM | POPC bilayer | CG-MetaD-M2 | 1 | 153 |
| 28 | TrkB TM | POPC bilayer | CG-MetaD-M2 | 1 | 120 |
| 29 | p75-C257A TM | POPC bilayer | CG-MetaD-M2 | 1 | 144 |
| 30 | TrkA TM | POPC bilayer | CG-MetaD-M3 | 1 | 113 |
| 31 | TrkB TM | POPC bilayer | CG-MetaD-M3 | 1 | 106 |
| 32 | p75-C257A TM | POPC bilayer | CG-MetaD-M3 | 1 | 125 |

[1] The tendency of the CG systems to explore TM arrangements beyond the NMR structures was examined by performing some additional test simulations. In the simulations presented in the main text, the TM helices were restrained in the equilibration phase while the micelle was being formed around them, and the restraints were lifted in the production phase, when an equilibrated DPC micelle had been formed (see Materials and Methods section). The effects of micelle equilibration were assessed by running some test simulations with Martini 2.2 where the TM helices were allowed to move unrestrained while the micelle was being formed around them, i.e. in the equilibration step (Systems 7-10). Comparison of the population density heatmaps acquired from these test simulations with the previous ones showed similar arrangements explored by the systems. Additional test simulations were performed using 24 mM ionic strength and a detergent-to-protein ratio of 20:1 (Systems 11,12), which are described in the experimental section of the TrkA NMR structure PDB file (PDB ID: 2N90). Again, the population density heatmaps were similar to those presented in Figure 2, suggesting that the results are robust and are not affected by these changes in simulation conditions.

**Table S2: Sequence-based prediction of the human TrkB TM domain.** Predictions of various bioinformatics webservers and databases are given for the TrkB sequence for the TM part (upper table) or for the secondary structure type (lower table). The prediction of each server for the TM sequence is highlighted in green in the upper table, while the predicted helical or strand regions are highlighted in green and orange, respectively, in the lower table. The dashed lines denote the beginning and the end of the region that was modeled as helical in the final model used for the molecular dynamics simulations.

| Prediction Source | TM predicted sequence |
|---|---|
| | 427 433 458 465 |
| UniProt | GREHLSVYAVVVIASVVGFCLLVMLFLLKLARHSKFGMK |
| ELM | GREHLSVYAVVVIASVVGFCLLVMLFLLKLARHSKFGMK |
| MEMSAT (PHYRE²) | GREHLSVYAVVVIASVVGFCLLVMLFLLKLARHSKFGMK |
| MEMSAT-SVM | GREHLSVYAVVVIASVVGFCLLVMLFLLKLARHSKFGMK |
| TMpred (in to out helix) | GREHLSVYAVVVIASVVGFCLLVMLFLLKLARHSKFGMK |
| TMpred (out to in helix) | GREHLSVYAVVVIASVVGFCLLVMLFLLKLARHSKFGMK |
| TMHMM | GREHLSVYAVVVIASVVGFCLLVMLFLLKLARHSKFGMK |
| PredictProtein | GREHLSVYAVVVIASVVGFCLLVMLFLLKLARHSKFGMK |

| Prediction Source | Strand-helix predicted sequence |
|---|---|
| Jpred4 | GREHLSVYAVVVIASVVGFCLLVMLFLLKLARHSKFGMK |
| PSIPRED | GREHLSVYAVVVIASVVGFCLLVMLFLLKLARHSKFGMK |
| NPSA-PRABI | GREHLSVYAVVVIASVVGFCLLVMLFLLKLARHSKFGMK |
| CFSSP | GREHLSVYAVVVIASVVGFCLLVMLFLLKLARHSKFGMK |

**Supplementary Data 1: Mapping between Martini coarse-grained beads and atoms for a DPC lipid molecule.** This mapping was used in the backward.py script [1] for conversion of the CG DPC micelle to atomistic representation.

```
[ molecule ]
DPC
[ martini ]
NC3 PO4 C1 C2 C3
[ mapping ]
charmm36
[ atoms ]
; Terminal head group (choline)
   1    N   NC3
   2   C13   NC3
   3  H13A   NC3
   4  H13B   NC3
   5  H13C   NC3
   6   C14   NC3
   7  H14A   NC3
   8  H14B   NC3
   9  H14C   NC3
  10   C15   NC3
  11  H15A   NC3
  12  H15B   NC3
  13  H15C   NC3
  14   C12   NC3 NC3 NC3 PO4
  15  H12A   NC3 NC3 NC3 PO4
  16  H12B   NC3 NC3 NC3 PO4
  17   C11   NC3 PO4
  18  H11A   NC3 PO4
  19  H11B   NC3 PO4
; Phosphate group
  20    P   PO4
  21   O13   PO4
  22   O14   PO4
  23   O11   PO4 PO4 C1
  24   O12   PO4 PO4 PO4 NC3
; Acyl chain
  25   C31   C1 PO4
  26   H1X   C1 PO4
  27   H1Y   C1 PO4
  28   C32   C1 C1 PO4
  29   H2X   C1 C1 PO4
  30   H2Y   C1 C1 PO4
  31   C33   C1 C1 C1 PO4
  32   H3X   C1 C1 C1 PO4
```

```
  33  H3Y   C1 C1 C1 PO4
  34  C34   C1
  35  H4X   C1
  36  H4Y   C1
  37  C35   C1 C1 C2
  38  H5X   C1 C1 C2
  39  H5Y   C1 C1 C2
  40  C36   C2 C1
  41  H6X   C2 C1
  42  H6Y   C2 C1
  43  C37   C2 C2 C1
  44  H7X   C2 C2 C1
  45  H7Y   C2 C2 C1
  46  C38   C2
  47  H8X   C2
  48  H8Y   C2
  49  C39   C2 C2 C3
  50  H9X   C2 C2 C3
  51  H9Y   C2 C2 C3
  52  C310   C3 C2
  53  H10X   C3 C2
  54  H10Y   C3 C2
  55  C311   C3 C3 C2
  56  H11X   C3 C3 C2
  57  H11Y   C3 C3 C2
  58  C312   C3
  59  H12X   C3
  60  H12Y   C3
  61  H12Z   C3
;;;making a choline group
[out]
C14 N C13 C12
H14A N C13 C12
H14B N C13 C12
H14C N C13 C12
[ chiral ]
C15 N C12 C13 C14
H15A N C12 C13 C14
H15B N C12 C13 C14
H15C N C12 C13 C14
```

**<u>Supplementary References</u>**


[1] T. A. Wassenaar, K. Pluhackova, R. A. Böckmann, S. J. Marrink, and D. P. Tieleman, "Going Backward: A Flexible Geometric Approach to Reverse Transformation from Coarse Grained to Atomistic Models," *J. Chem. Theory Comput.*, vol. 10, no. 2, pp. 676–690, Feb. 2014.
